# A transient modified mRNA encoding Myc and Cyclin T1 induces cardiac regeneration and improves cardiac function after myocardial injury

**DOI:** 10.1101/2023.08.02.551469

**Authors:** Aleksandra Boikova, Gregory A. Quaife-Ryan, Christopher A.P. Batho, Elsa Lawrence, Harley Robinson, Camilla Ascanelli, Karin Jennbacken, Qing-Dong Wang, Kenny M. Hansson, Adam Seaton, Victoria Rodriguez Noci, Megan Bywater, Jasmin Straube, Kamil A. Sokolowski, Brian W.C. Tse, Thomas Krieg, Ana Vujic, Enzo R. Porrello, Sanjay Sinha, James E. Hudson, Catherine H. Wilson

## Abstract

Cardiac injury, such as myocardial infarction (MI), results in permanent loss of cardiomyocytes and in many cases heart failure. Transgenic expression of the pro-proliferative transcription factor Myc and Cyclin T1 can drive substantial adult cardiomyocyte proliferation to replace lost cardiomyocytes. Herein, we show that Myc and Cyclin T1 induced cardiomyocyte proliferation leads to myocardial repair and functional (long-term) recovery post-MI in mice. To provide a more translational approach, we developed modified mRNA (modRNA) encoding Myc-Ccnt1 as a transient and non-integrating strategy for regeneration. One dose of Myc-Ccnt1 modRNA is sufficient to transiently drives cardiomyocyte proliferation in human pluripotent stem cell-derived cardiomyocytes and a mouse MI model, where it leads to better heart function. Using single nuclei sequencing and proteomics, we show this was functionally mediated by transcriptional activation of cell-cycle regulating genes, which ultimately results in mitosis and cytokinesis of cardiomyocytes. Collectively, these findings indicate that Myc-Ccnt1 modRNA has the potential to be an effective regenerative therapeutic.

## Introduction

Cardiovascular disease is the leading cause of death worldwide, causing >18 million deaths annually. Of these deaths, around 85% are attributed to heart attack and stroke^1^. Following heart attack or myocardial infarction (MI), cardiomyocytes perish and are never replaced. The permanent loss of contractile cells leads to deterioration of heart function and ultimately heart failure where the 5-year mortality rate is around 50%^2^. Current therapies for ischaemic heart failure include urgent revascularisation and relieving the symptoms, by reducing blood pressure and blood volume. Artificial ventricular assist devices are available but are not yet suitable for long-term use due to complications with infection and thrombosis. None of these options can reverse the muscle cell loss and the degeneration of the heart tissue. The only effective treatment for end-stage heart failure is heart transplantation, but the global shortage of hearts for organ donation limits transplants to only thousands per year and requires patients to undergo aggressive immunosuppressive regimens. Driving the heart to endogenously generate new cardiomyocytes is one of the most attractive treatment strategies to recover contractile function and reverse heart failure. Hence, there has been intensive interest in uncovering the molecular programs that control cardiomyocyte proliferative capacity and utilising these factors to augment adult cardiomyocyte turnover for heart regeneration. Several mitogens potentiate adult cardiomyocyte proliferation but despite decades of concerted efforts, there are no clinically approved regenerative therapeutics for the adult heart following injury.

We recently showed that transgenic expression of Myc and increasing levels of the positive transcription elongation factor (P-TEFb) complex drives substantial adult murine cardiomyocyte proliferation^3, 4^. Myc is a proliferation activating transcription factor that serves as a pivotal, non-redundant instructor of tissue regeneration following injury^5^. Myc expression in normal cells is stringently controlled. Following damage in a regenerative tissue, local mitogenic stimulation leads to a transient increase in short-lived Myc protein followed by a rapid burst of proliferation^5^. In cells with low proliferative capacity, such as adult cardiomyocytes, we have determined that ectopic expression of Myc alone cannot drive successful cell division, due to the low levels of P-TEFb within the cell^3, 4, 6, 7^. P-TEFb is a protein complex made up of CDK9 and Cyclin T1 (encoded by *Ccnt1*) that, following stimulation, is recruited to and phosphorylates RNA Pol II to enable productive transcription^8^. Cyclin T1 is the key factor regulating P-TEFb levels in cardiomyocytes^3, 6, 7^. Combined overexpression of Myc and Cyclin T1 in cardiomyocytes leads to efficient Myc-driven transcription that can drive juvenile and adult mouse cardiomyocyte proliferation *in vivo*^3, 4^ leading us to speculate whether this genetic combination could be explored for therapeutic benefits after cardiac injury.

The cardiomyocyte proliferative response to Myc and Cyclin T1 in the mouse heart is potent. Sustained whole heart expression in 15-day old mice leads to functional decline and death over 4 days^3^. In contrast, transient Myc expression in adult mice with a limited number of cardiomyocytes ectopically expressing Cyclin T1 (30%) is well-tolerated^3, 4^. Here we use a genetically modified mouse model of combined ectopic expression of Myc and Cyclin T1 together with LAD coronary artery ligation to show that an efficient therapeutic response is achieved when, Myc expression is localised to the damaged area. Next, we hypothesise that for a regenerative therapeutic to be successful, careful titration of the treatment dose, timing, location, and duration of Myc and Cyclin T1 are required. To test this, we utilised a modified mRNA (modRNA) encoding *Myc* and *Ccnt1* to facilitate localised and transient expression as a prospective cardiac regenerative therapeutic. Our results demonstrate that delivery of *Myc* and *Ccnt1* modRNA drives transient expression of Myc and Cyclin T1 protein and activation of downstream Myc programmes that provokes cardiomyocyte proliferation and functional repair. Establishing that localised and transient delivery cardiac regenerative genes is an important therapeutic strategy for cardiac regeneration.

## Results

### MycER^T2^ and Cyclin T1 expression drives enhanced proliferation of cardiomyocytes following MI

We employed the cardiac specific, tamoxifen controlled *Myh6-Cre;Gt(ROSA)26Sor^tm3(CAG–MYC/ERT2)Gev^* (*CMER*) mice to establish whether whole heart transient expression of MycER^T2^ combined with Cyclin T1 could drive cardiac repair in adult mice. At 4 weeks of age *Myh6-Cre;CMER* and *CMER* Cre negative mice were systemically injected with an adeno-associated virus encoding *Ccnt1* (*AAV9-cTnT-Ccnt1*). At 8 weeks, mice underwent left anterior descending coronary artery ligation (LAD) to induce MI. Mice were treated with tamoxifen, to transiently activate MycER^T2^, or oil (vehicle control) by intraperitoneal injection at day 1 and day 2 following LAD and collected at 72 hours post MI for histological and transcriptional analysis (Figure 1A). Similar to our previous finding^3,4^, we observed Ki67 (general cell-cycle; control (cre + oil) 0.2%, Myc + Cyclin T1 (cre + tam) 25.1%), P-H3 (mitotic; control (cre + oil) 0.5%, Myc + Cyclin T1 (cre + tam) 6.9%) and Aurora B kinase (interphase to cytokinesis; control (cre + oil) 1.4%, Myc + Cyclin T1 (cre + tam) 4.4%) positive cardiomyocytes in Myc and Cyclin T1 expressing hearts, indicating efficient cell cycle entry and progression (Figure 1B and C, Supplemental Figure 1A and B). The frequency of Ki67, P-H3 and Aurora B kinase positive cardiomyocytes was significantly higher in the border zone (BZ; 25.1%, 6.9% and 4.4% respectively for Myc + Cyclin T1 (cre + tam)), the area adjacent to the infarcted myocardium, when compared to the remote zone (RZ; 6.2%, 2.9% and 0.6% respectively for Myc + Cyclin T1 (cre + tam)), the myocardium located distal from the infarcted region. Next, we used Terminal deoxynucleotidyl transferase dUTP nick end labelling (TUNEL) and cleaved caspase 3 staining to quantitatively determine the level of apoptotic cardiomyocytes and cell death in the heart. We show that elevated Myc and Cyclin T1 activity alone does not lead to cardiomyocyte cell death when compared to control hearts (Figure 1D). Thus, the enhanced cardiomyocyte proliferation in the border zone of the infarct, specifically in the Myc and Cyclin T1 heart, is a response to the induced MI.

**Figure 1:**
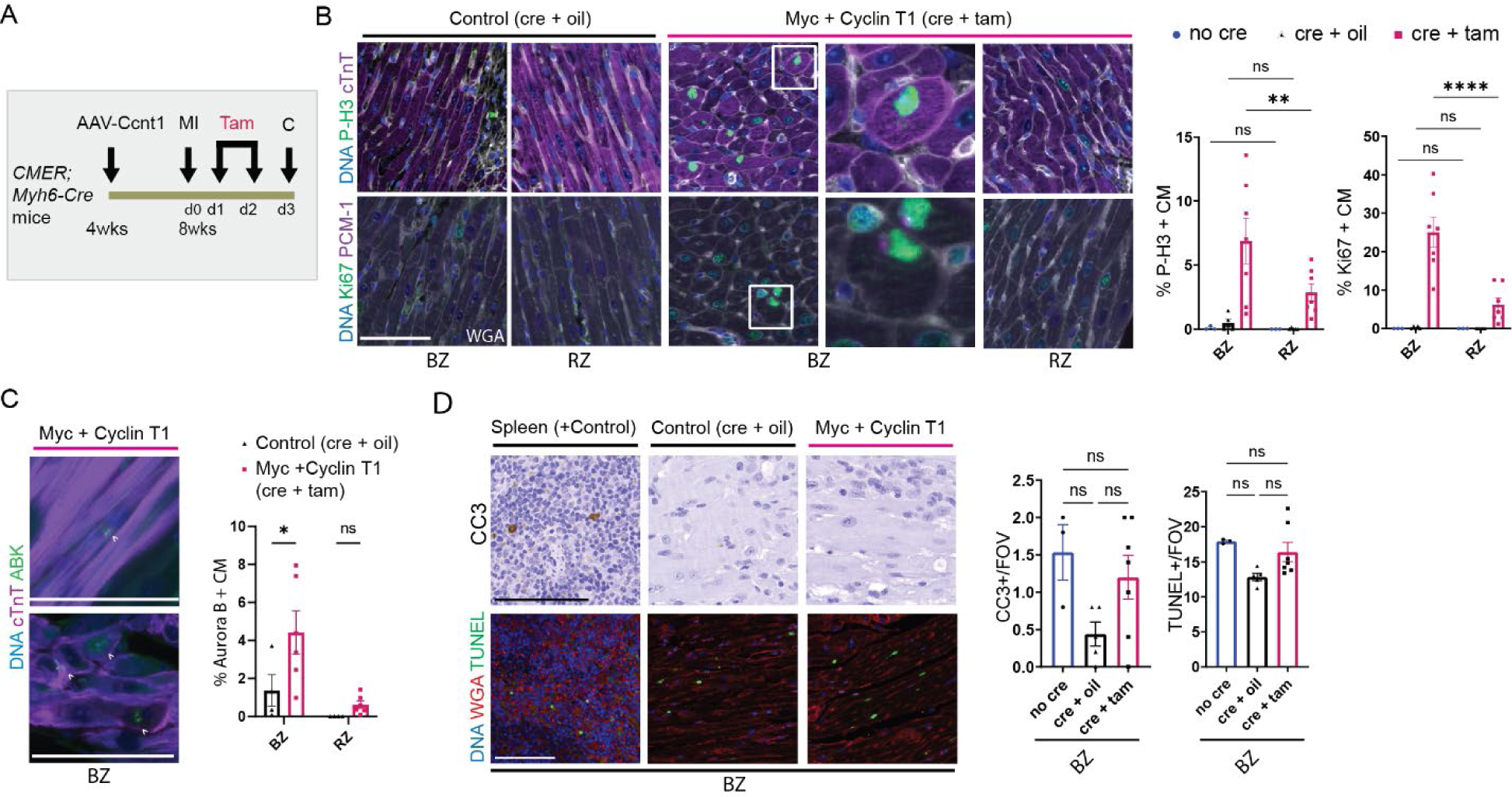
MycER^T2^ and Cyclin T1 expression lead to increased cell cycle marker expression in cardiomyocytes in vivo. (A) An experimental schematic to determine the proliferative potential of MycER^T2^ and Cyclin T1 overexpression adult mice. (B) Immunofluorescence staining and quantification of P-H3 or Ki67 (green), cardiac troponin or PCM-1 (purple), wheat germ agglutinin (WGA, white) and DNA (Blue) in the border or remote zone of MycER^T2^ and Cyclin T1 (Cyclin T1 AAV and cre + tam, n = 7) and control (Cyclin T1 AAV and either oil treated (cre + oil, n = 5) or cre negative (no cre, n = 3)) hearts 48 hours following activation of MycER^T2^ by tamoxifen treatment. Two way-ANOVA with Šídák’s multiple comparisons test BZ to RZ; P-H3 p = 0.0021 (**), Ki67 p < 0.0001 (****). (C) Immunofluorescence staining and quantification of Aurora B Kinase (green), cardiac troponin (purple) and DNA (blue) in the border and remote zone of MycER^T2^ and Cyclin T1 (cre + tam, n = 6) and control (either oil treated (cre + oil, n = 4)) hearts 48 hours following activation of MycER^T2^ by tamoxifen treatment. White arrows point to stages of positivity, including symmetrical midbody (top image). Two way-ANOVA with Šídák’s multiple comparisons test cre + tam vs cre + oil in the BZ p = 0.0287 (*). (D) Immunofluorescence and immunohistochemical staining and quantification of Cleaved caspase 3 (CC3) and TUNEL (green), WGA (red) and DNA (blue) in the border zone of MycER^T2^ and Cyclin T1 (cre + tam, n = 7) and control (either oil treated (cre + oil, n = 5) or cre negative (no cre, n = 3)) hearts 48 hours following activation of MycER^T2^ by tamoxifen treatment. Spleen is shown as a positive control for staining. One way-ANOVA not significant (ns). No staining was observed in the RZ. All scale bars are 100 µm. Mean and SEM displayed.

### Single nuclei RNA sequencing defines expression changes in proliferative cardiomyocytes

We performed single nuclei RNA sequencing (snRNAseq) on hearts collected from adult *Myh6-Cre;CMER* (n = 3) and *CMER* cre negative (n = 3) mice that were systemically injected with AAV9-*Ccnt1* at 4 weeks of age and that underwent MI at 8 weeks, followed by MycER^T2^ activation for 48 hours (Figure 1A). After quality control and doublet removal (Supplemental Figure 2A) and using previously published sets of markers for snRNAseq of whole mouse heart^9^ we identified 12 transcriptionally distinct cell clusters (Figure 2A to C). In addition to adult injured hearts, we performed snRNAseq of hearts collected from AAV9-*Ccnt1* treated adult mice that had not undergone MI (adult no-MI, *Myh6-Cre;CMER,* n=4 and *CMER* cre negative, n = 2) and 15 day juvenile mice (*Myh6-Cre;CMER*, n = 3 and *CMER* cre negative, n = 3) both 48 hours post tamoxifen injection (Supplement Figure 2B and C). We have previously shown that 15-day juvenile mice express endogenously high levels of cardiac Cyclin T1 and are extensively permissive to Myc-driven transcription, with the activation of ectopic Myc leading to widespread cardiomyocyte proliferation^3^. We found the same cell types and saw a similar distribution of populations (Supplemental Figure 2B to G) in all 3 data sets. MycER^T2^ and Cyclin T1 are exclusively expressed in cardiomyocytes therefore, to allow cross-sample comparisons we integrated and analysed the cardiomyocyte populations (Figure 2D). We observed specific Myc target gene expression in MycER^T2^ + Cyclin T1 expressing cardiomyocytes (Supplemental Figure 3A), and differential gene expression analysis indicated a large overlap in upregulated and downregulated genes over the 3 data sets (Figure 2E and Supplemental Figure 3B); juvenile and adult MI datasets showing the largest overlap. Gene set enrichment determined that MycER^T2^ + Cyclin T1 expressing cardiomyocytes expressed transcripts overlapping with hallmark genes lists such as Myc targets, G2M targets and E2F targets, while down regulated genes overlapped with genes involved in myogenesis (Figure 2F and G), indicating that the transcriptional signatures of cardiomyocytes following ectopic MycER^T2^ and Cyclin T1 are reflective of cells with elevated Myc transcriptional activity and progressing through mitosis.

**Figure 2:**
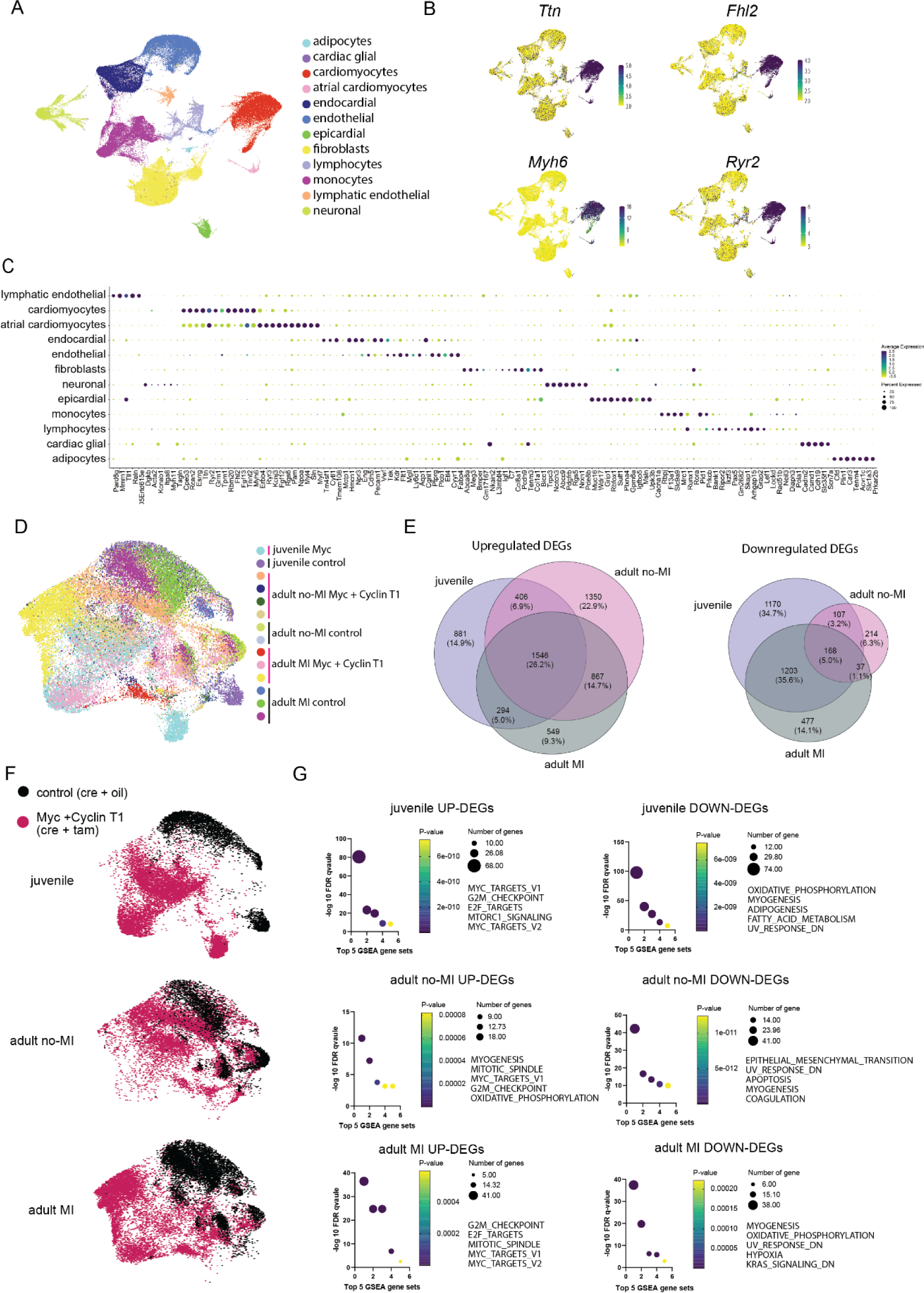
Cardiomyocytes expressing MycER^T2^ and Cyclin T1 display transcriptional markers of cell cycle and cytokinesis. (A) UMAP representation of 12 major cell types identified from 121394 recovered nuclei in the adult MI heart dataset. (B) UMAP plots showing expression of ventricular cardiomyocyte specific genes *Ttn*, *Fhl2*, *Myh6*, *Ryr2*. (C) Dot plot showing expression of canonical marker gene panel for 12 cell types. (D) UMAP plot of subset and integrated cardiomyocytes showing the overlap between 12 samples from 3 datasets (juvenile, adult no-MI, and adult MI). We recovered 46467 total cardiomyocyte nuclei. (E) Venn diagrams showing overlap between differentially expressed upregulated and downregulated genes between MycER^T2^ + Ccnt1 and control conditions in 3 datasets (DESeq2, p_adj < 0.01). (F) UMAP plot showing overlap between Myc + Ccnt1 and control conditions (juvenile, adult no-MI, and adult MI). (G) Top 5 enriched GSEA Hallmark gene sets for top 400 upregulated and bottom 400 downregulated differentially expressed genes from MSigDB.

Cardiomyocyte sub-cluster analysis, irrespective of Myc activation, identified 4 distinct cell states with unique transcriptional expression (Figure 3A). Cardiomyocyte cluster 1 (CM1) was predominant in the control samples comprising 67%, 52% and 71% of cardiomyocytes in the juvenile, adult no-MI and adult MI controls respectively (Supplemental Figure 3C). CM1 was characterised by expression of mature ventricular cardiomyocyte markers such as Ryr2, Ttn and Myh6 and biological processes involved in cardiac muscle contraction (Figure 3B and Supplemental Figure 3D). Cardiomyocyte cluster 2 (CM2) was enriched in samples overexpressing MycER^T2^ and Cyclin T1 accounting for 59%, 57% and 36% of cardiomyocytes in Myc expressing juvenile, MI and no-MI respectively and 2%, 44% and 19% in controls (juvenile, MI and no-MI respectively). CM2 displayed markers such as Xirp2, Creb5, Grk5 involved in cardiac conduction and cellular cardiomyocyte stress response^10, 11^. Cardiomyocyte cluster 3 (CM3) was the most distinct and dynamic population, predominantly observed in MycER^T2^ expressing samples from adult MI (14% vs. > 0.1% in controls) and juvenile mice (21% vs. > 0.1% in controls). CM3 was characterised by expression of cell cycle genes such as Top2a, Cenpp, Mki67 and Ccnd3 and biological processes involved in cell cycle. Cardiomyocyte cluster 4 (CM4) was identified predominantly in juvenile hearts accounting for 18% of juvenile cardiomyocytes expressing Myc and 31% of juvenile control cardiomyocytes. CM4 was characterised by markers associated metabolism such as Mb, Cox6c, Atp5, Fabp3 and biological processes involved in cellular respiration (Figure 3B and Supplemental Figure 3B and C). Guided by the subcluster markers, we performed cell cycle scoring analysis which indicated that CM cluster 3 was highly enriched for S-phase (e.g., Top2a and Mki67), G2M (e.g., Cdk1 and Ccnb2), and cytokinesis markers (Anln and Cenpa) (Figure 3C and D). In agreement with the histological data (Figure 1B), the transcriptional data indicated an increase in cardiomyocytes with cell cycle markers following Myc and Cyclin T1 expression post-MI compared to an undamaged setting (Figure 3C, adult MI vs adult no-MI).

**Figure 3:**
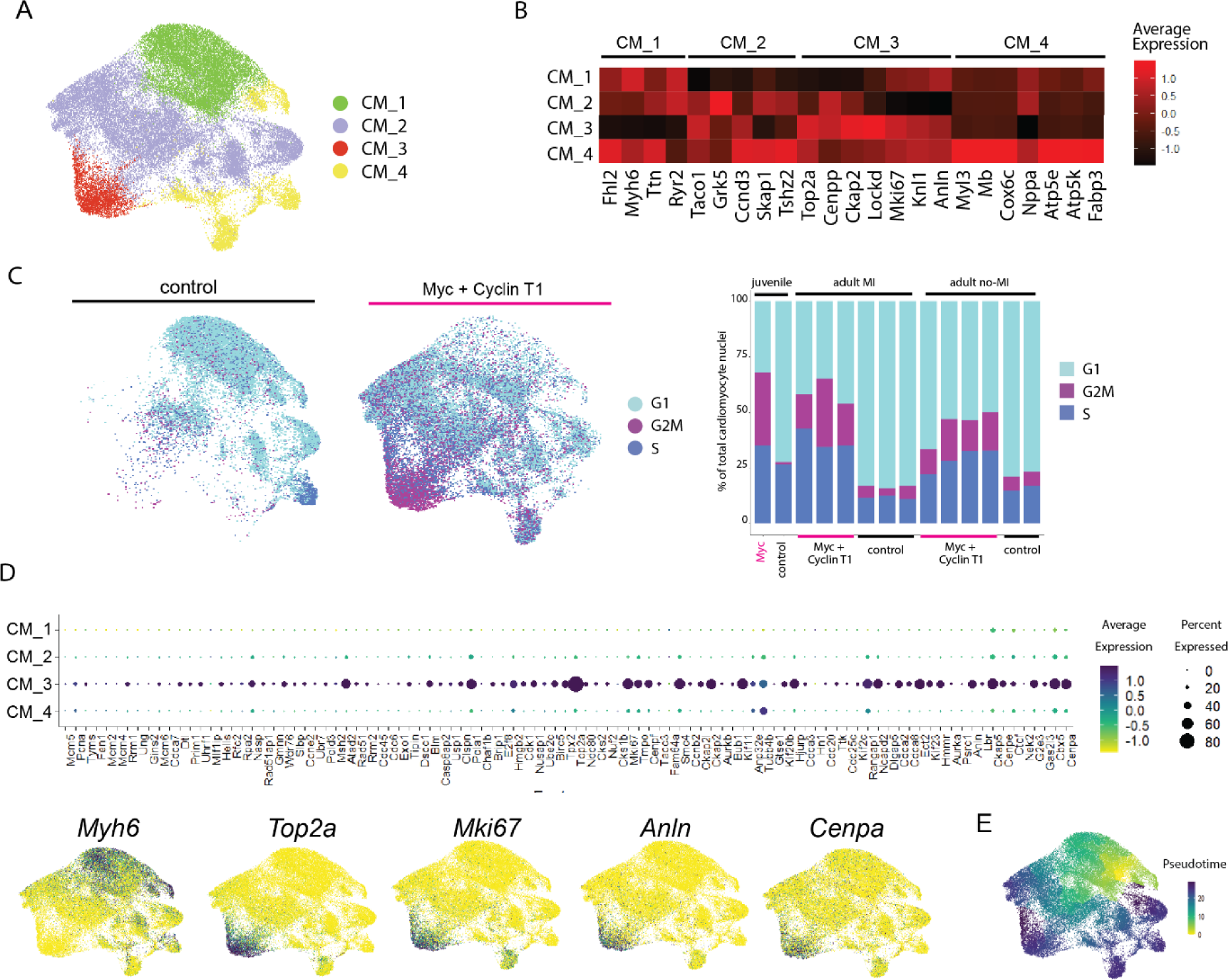
Cardiomyocytes expressing MycER^T2^ and Cyclin T1 show Myc driven cell cycle re-entry and progression. (A) UMAP plot of 4 cardiomyocyte subclusters CM1, CM2, CM3, CM4. (B) Average expression heatmap of selected top differentially expressed subcluster marker genes per subcluster (Wilcox test, positive log2FC > 0.25, p_adj < 0.01). (C) UMAP plot of cardiomyocyte cell cycle phase scoring (left) split by genotype group control or MycER^T2^ + Ccnt1 and control. Histogram (right) of cell cycle phase scoring nuclei as % of total nuclei per biological replicate sample in juvenile, adult MI and adult no-MI datasets. (D) Dot plot (top) of canonical cell cycle S-phase, G2M and cytokinesis gene panel per cardiomyocyte subcluster. UMAP feature plots (bottom) for cardiomyocyte marker *Myh6* and cell cycle markers *Top2a*, *Mki67*, *Anln* and *Cenpa* for cardiomyocyte subclusters. (E) Monocle 3 pseudotime cell trajectory originating from the CM1 subcluster control nuclei enriched node.

Pseudotemporal cell trajectory analysis indicated that the cell state changed from CM1 and progressed though CM2 to CM3 clusters (Figure 3E), suggesting cardiomyocytes passage through a dynamic biological path to division. Further analysis of subclusters CM1 to 3 demonstrated Myc-driven activation of mTORC1 signalling, glycolysis and oxidative phosphorylation in cardiomyocytes, indicating that Myc plays a central role instructing growth and metabolic remodelling during cardiomyocyte cell cycle (Supplemental Figure 3E). Emergence of CM3, 48 hours following MycER^T2^ activation suggests that this new CM cluster is derived from another cardiomyocyte population that may have undergone transcriptional remodelling.

### Localised MycER^T2^ and Cyclin T1 drives functional recovery of the heart following MI

To determine whether cardiomyocyte specific Myc and Cyclin T1 expression could provide therapeutic benefit following cardiac injury we generated a cohort of adult *Myh6-Cre;CMER* and *CMER* Cre negative mice for echocardiogram analysis (Figure 1A). Mice were systemically injected with AAV9-*Ccnt1* at 4 weeks of age and at adulthood MI was performed. MycER^T2^ was activated with 2 injections of tamoxifen on day 1 and 2 following MI. No post-MI echocardiogram analysis was possible because at 3 to 7 days post-MycER^T2^ activation, 89% (8/9) mice that carried *Myh6-Cre;CMER* were culled moribund or found dead whereas no (0/8) *CMER* Cre negative were culled due to deteriorating clinical signs. This is in stark contrast to the 100% survival of *Cre;CMER + AAV Cyclin T1* + Tam treated mice at 28 days following MycER^T2^ activation which had not undergone MI surgery^3^. Histological analysis of available hearts indicated high numbers of proliferative cardiomyocytes (Supplemental Figure 4A) which we hypothesize caused a detrimental reduction in heart function, however, cause of death was not possible to confirm.

To reduce the overall number of cells expressing MycER^T2^, we established a method of direct myocardial injection of modRNA encoding Cre to localise expression of MycER^T2^ to the cardiomyocytes surrounding the infarct site. *Gt(ROSA)26Sor^tm14(CAG-tdTomato)Hze^* (*tdTomato*) reporter mice were infarcted and intramyocardially injected with 100 µg Cre modRNA during surgery, immediately post-MI. Cre-mediated recombination could be observed in ∼9% of the left ventricle (Figure 4A), similar to previous reports^12^. Non-heart Cre mediated recombination was extremely rare in the kidney, thymus and lung, and was observed in 0.14% and 0.27% of cells in the liver and spleen respectively. (Supplemental Figure 4B). To establish the kinetics of modRNA expression, adult CD1 mice were infarcted and intramyocardially injected with 100 µg FLuc modRNA. FLuc bioluminescence exhibited a localised and “pulse-like” expression, with expression peaking early in the heart and rapidly declining with a 5.5-fold reduction in expression from 8 to 48 hours post injection (Figure 4B) similar to previously published kinetics of modRNAs^13^.

**Figure 4:**
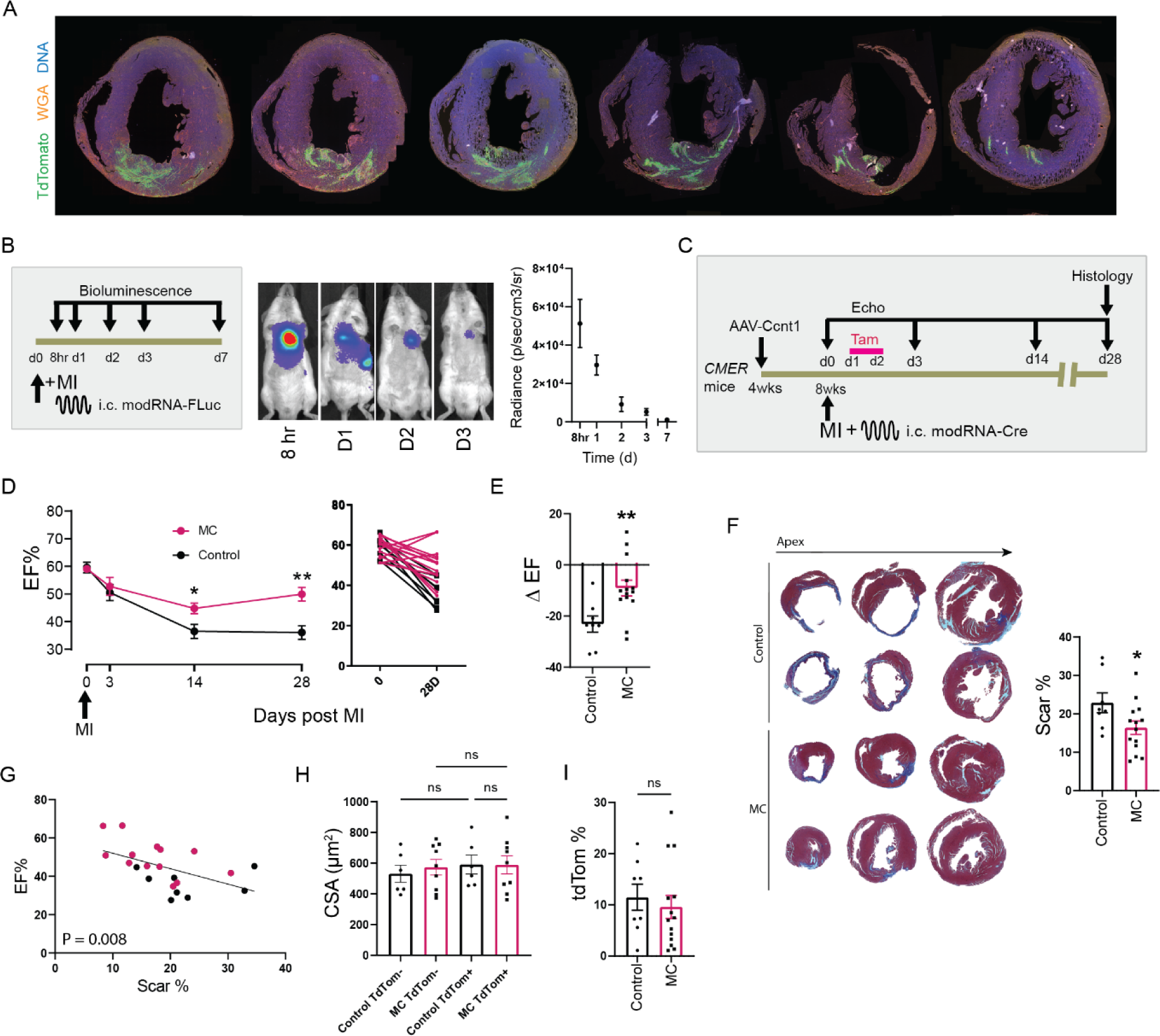
Localised MycER^T2^ and Cyclin T1 elicit cardiac benefit following MI in mice. (A) Immunofluorescence staining through a representative *tdTomato* reporter mouse heart where *tdTomato* (green), WGA (orange) and DNA (blue) are visualised 5 days following intramyocardial injection with 100 µg Cre modRNA during surgery, immediately post-MI. (B) An experimental schematic to study intramyocardially injected FLuc modRNA (FLuc) assessed with bioluminescence. IVIS imaging and quantification of mice following intramyocardial injection of FLuc at 8 hours and 1, 2, 3 and 7 days. n = 4 (8hr), n = 3 (D1), n = 4 (D2), n = 6 (D3) and n = 5 (D7). (C) An experimental schematic to study the cardioregenerative potential of localised and transient MycER^T2^ and Cyclin T1 (MC) in 8-week-old mice. Cardiac function was assessed with echocardiography on days 0, 3, 14 and 28 post MI. Mice were sacrificed at day 28 for histology. n = 8 control and n = 14 MC. (D) Mean percent ejection fraction (LV long-axis) of MC and control mice over 28 days post MI. Individual mice shown at 0 and 28 days. Unpaired t-test at 14 days p = 0.0016 and at 28 days p = 0.0193. (E) Difference in %EF between MC and control mice (ΔEF% calculated by D0 (pre-MI) – D28 (post-MI) EF%) (p = 0.0075, unpaired t-test). (F) Masson’s Trichrome staining of representative hearts from MC and control mice at 28 days post MI. Scar area/total heart area was quantified and analysed using an unpaired t-test (P = 0.0434). (G) Spearman rank correlation between Scar % and EF % from MC (pink) and control (black) mice at 28 days. P = 0.0080 Slope = Y = −0.8048*X + 60.10. (H) Quantification of cardiomyocyte cross sectional area (CSA) (n.s.- not significant). (I) Quantification of the percent immunofluorescence positivity of *tdTomato* in hearts from MC and control mice at 28 days post MI, (n.s.- not significant). Mean and SEM shown.

We sought to identify whether restricted, localised overexpression of MycER^T2^ (and Cyclin T1) could enable regeneration of the adult murine heart after MI. CMER;*tdTomato or tdTomato* mice were systemically injected AAV9-*Ccnt1* at 4 weeks of age and at adulthood were infarcted and intramyocardially injected with 100 µg of Cre modRNA. Mice were recovered from MI surgery and on day 1 and day 2 post-surgery mice were treated with tamoxifen injections to activate MycER^T2^ (Figure 4C). We assessed cardiac function before and post MI at 3, 14 and 28 days and MycER^T2^ expressing mice demonstrated greater cardiac function at 28 days post MI compared to control (Figure 4D, Supplemental figure 4C). The change in left ventricular ejection fraction (ΔEF%) at day 28 post-MI for control and Myc + Cyclin T1 (MC) were −23±3 and −9±3 %, respectively (Figure 4E). At day 28 post-MI we observed a reduced scar area in MycER^T2^ + Cyclin T1 expressing mice compared to controls (Figure 4F), and the percent scar area negatively correlated with LVEF% across the whole cohort (Figure 4G). Cardiomyocyte surface area did not differ between MycER^T2^ expressing mice and controls (Figure 4H). Histological analysis of tdTomato indicated that overall, 10% of cardiac tissue had been recombined across the cohorts and there was no difference in percent recombined area between MycER^T2^ expressing mice and the controls (Figure 4I). Rare Tdtomato positive-Cre recombined cells were observed in spleen and liver in similar numbers to control mice (Supplemental Figure 4D).

### A modified mRNA prototypical therapeutic encoding Myc and Cyclin T1 drives transient expression and cardiomyocyte proliferation *in vitro*

To develop a therapeutic based on Myc and Cyclin T1 expression, mouse and human cMyc and Cyclin T1 ORF sequences were utilised to construct templates to synthesize a single modRNA encoding Myc and Ccnt1 separated by a T2A sequence (M-C) (Supplemental data, Figure 5A). We observed no significant differences between the function of human and mouse M-C proteins (Supplemental Figure 5A and B) in human and mouse cells *in vitro* therefore, for all other experiments we used mouse M-C modRNA.

**Figure 5:**
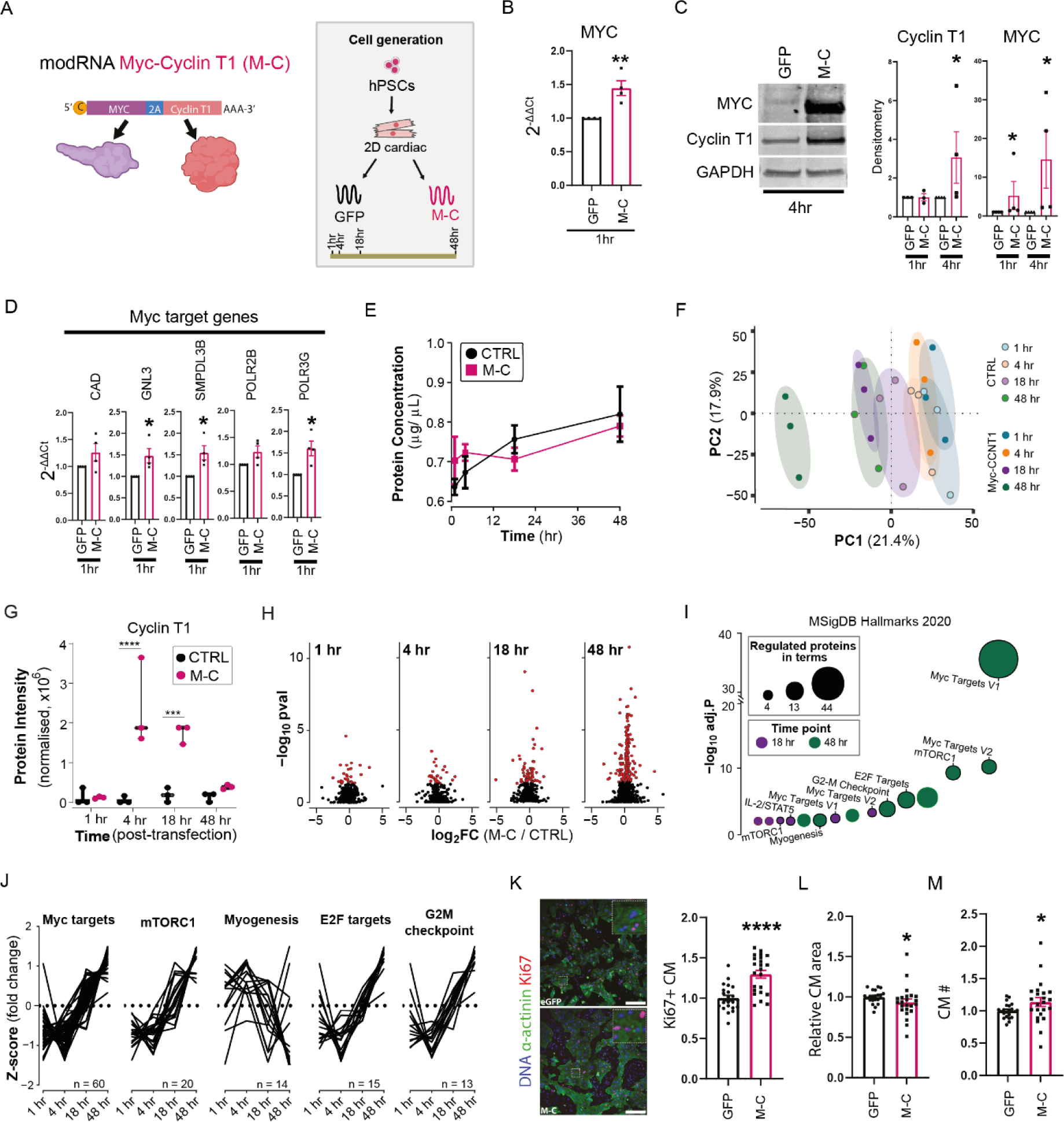
Myc and Cyclin T1 potentiates cardiomyocyte proliferation *in vitro*. (A) Illustration of Myc-T2A-Ccnt1 modRNA (M-C) and experimental workflow. (B) qRT-PCR analysis of Myc-Cyclin T1 (relative percent expression to HPRT) and Myc RNA expression in 2D cardiomyocytes at 1-hour post transfection (p = 0.006, n = 4, unpaired t-test). (C) Immunoblotting and quantification of Myc and Cyclin T1 protein levels at 1- and 4-hours post (M-C) modRNA treatment in 2D cardiomyocytes (n = 4, p = 0.029, Mann-Whitney test). (D) qRT-PCR analysis of downstream Myc responsive genes following M-C treatment, including SMPDL3B (p = 0.016), GNL3 (0.028), POLR3G (0.012). n = 4, unpaired t-test). (E) Concentration of protein collected from cultured cardiac cells at point of harvest. Mean and standard deviation denoted, (n = 3 for each condition). (F) PCA plot of each proteomics sample. (G) Cyclin T1 intensity measured by LC-MS/MS at each time point. 2-way ANOVA with Benjamini-Hochberg correction used for statistical evaluation between M-C and control (CTRL) conditions. (H) Volcano plots for all proteomic changes at each time points. Red values exceed the statistical threshold of p < 0.05 by Proteome Discoverer (2.3) analysis. (I) Gene set enrichment analysis of proteins significantly altered at 18 and 48 hr post-transfection. MSigDB Hallmarks 2020 terms were used to identify terms, where log transformed adjusted p-values (y-axis) and number of proteins identified per term (point size) denoted here. (J) Temporal trends of proteins belonging to the enriched pathways. Fold change data expressed as z-score scaled. (K) Immunofluorescence and quantification of Ki67 staining in GFP and M-C modRNA treated 2D cardiomyocyte cultures, scale bar 100 µm (p = <0.0001%, Mann-Whitney test). (L) Quantification of relative cardiomyocyte (CM) area (p = 0.0146, Mann-Whitney test) in 2D cardiomyocytes 48 hours post M-C transfection. (M) Quantification of cardiomyocyte number (p = 0.035, Mann-Whitney test) in 2D cardiomyocytes 48 hours post M-C transfection.

Using human embryonic stem cell-derived cardiomyocytes (hESC-CM) we confirmed expression of Myc and Cyclin T1 from M-C modRNA at the transcript and protein level following transfection at 1 and 4 hours (Figure 5B and C). Myc protein expression was diminished at 18 and 48 hours (Supplemental Figure 5C), suggesting ectopic Myc has a short protein half-life, similar to endogenous Myc^14, 15^. Treatment with a proteasome inhibitor increased Myc expression, indicating Myc stability was dependent on protein turnover (Supplemental Figure 5D). Downstream Myc-responsive genes (SMPDL3B, GNL3 and POLR3G) were transcriptionally upregulated as early as 1 hour post treatment with M-C (Figure 5D). We monitored the proteomic response by LC-MS/MS over time at 1-, 4-, 18- and 48-hours post-transfection of M-C modRNA (Figures 5E to J). Whilst the protein abundance of Myc and Cyclin T1 was transient and depleted by 48hrs (Figure 5G), M-C treatment led to a change in protein expression (Figure 5H) and translated Myc target genes were increased at 18- and 48-hours post transfection (Figure 5I). In addition to direct Myc target gene expression, we observed increased mTORC1, E2F targets and G2M checkpoint signatures (Figure 5J). We next established whether M-C treatment could drive cardiomyocyte proliferation, and at 48 hours post M-C modRNA treatment we saw significantly increased hESC-CM Ki67-positive and BrdU-positive hESC-derived cardiomyocytes (Figure 5K, Supplemental Figure 5A), with reduced cardiomyocyte cross-sectional area (Figure 5L) and increased cardiomyocyte cell number (Figure 5M). These data support M-C modRNA as a promising candidate to stimulate expression of Myc target genes and promote human cardiomyocyte proliferation.

### Transient Myc and Cyclin T1 expression result in cardiac benefit following myocardial infarction in mice

To assess M-C modRNA protein expression after MI in vivo adult CD1 mice were infarcted and intramyocardially injected with 100 µg of M-C, or GFP modRNA during surgery, immediately post-MI (Figure 6A). We confirmed GFP, Myc and Cyclin T1 protein expression in cardiomyocytes at 1- and 4-hours post injection (Figure 6B). Similar to hESC-CMs, Myc and Cyclin T1 protein expression was short-lived, and we did not observe protein expression at 24 hours in heart tissue, whilst GFP expression was maintained for at least 48 hours (Supplemental Figure 6A).

**Figure 6:**
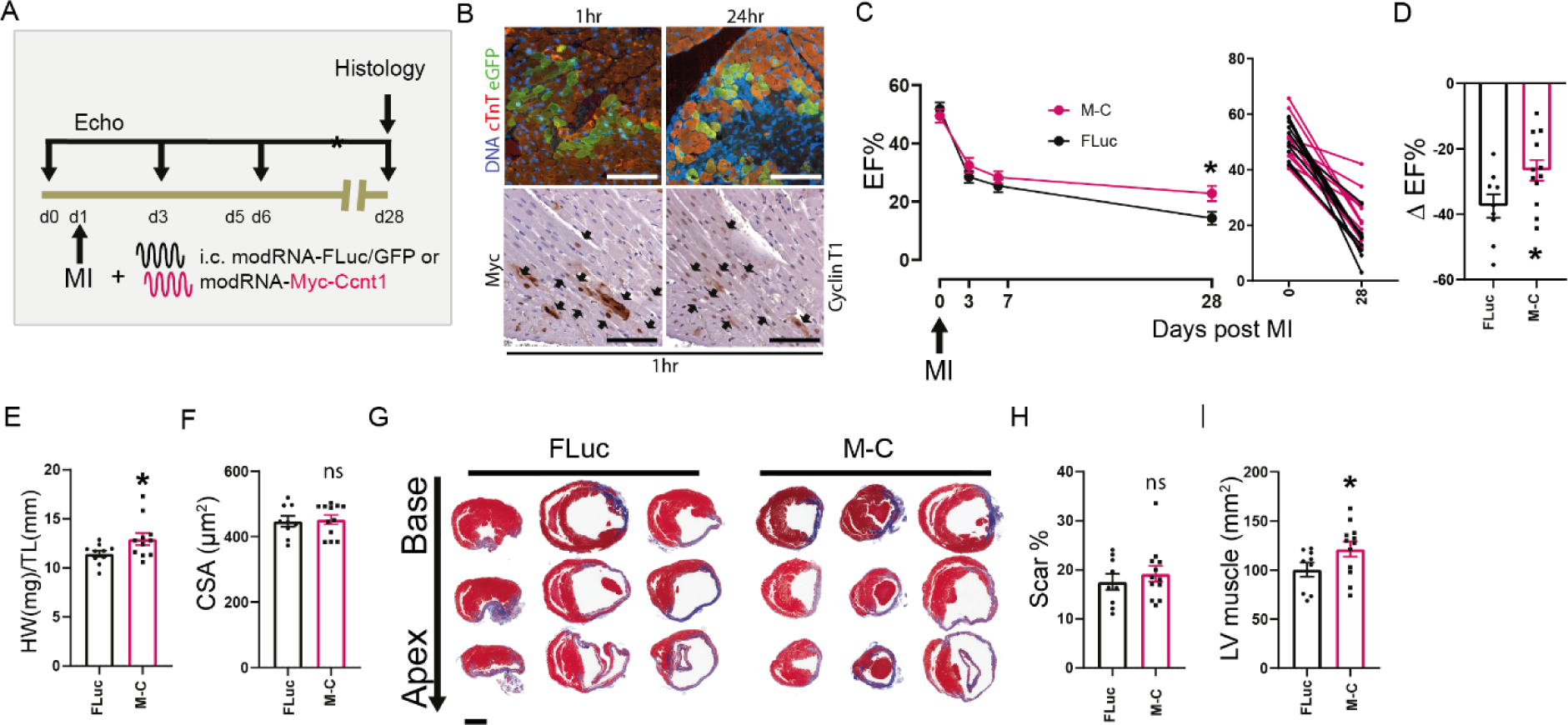
Assessment of cardiac function following M-C injection post MI. (A) An experimental schematic to study the cardioregenerative potential of M-C in 8-week old mice. Cardiac function was assessed with echocardiography on days 0, 3, 6 and 28 post MI. Mice were sacrificed at day 28 for histology. n = 9 FLuc and n = 12 M-C. (B) Immunofluorescence and Immunohistochemical staining of GFP (green), cardiac troponin (cTnT, red) DNA (blue), Myc and Cyclin T1 at 1- or 24-hour post modRNA injection. GFP protein expression is maintained at 24 hours, Myc and Cyclin T1 expression is absent by 24 hours. n = 3, scale bars 100 µm. (C) Mean ejection fraction % (left) of M-C and FLuc injected mice over 28 days post MI. Individual mice shown at 0 and 28 days (middle). Ejection fraction % (right) at 28 days, Unpaired t-test at 28 days p = 0.0301. (D) Difference in %EF (ΔEF% calculated by D0 (pre-MI) – D28 (post-MI) EF%). Unpaired t-test p = 0.0339. (E) Heart weight (mg)/Tibia length (mm) (HW/TL) of FLuc and M-C injected mice 28 days post MI and injection, Unpaired T-test, p= 0.048. (F) Quantification of cardiomyocyte cross sectional area (CSA) (n.s.- not significant). (G) Masson’s Trichrome staining of representative hearts from M-C and FLuc injected mice at 28 days post MI. (H) Quantification of scar area/total heart area (n.s.- not significant). (I) Left ventricular (LV) muscle area of FLuc and M-C injected mice 28 days post MI and injection (p = 0.0491, Mann-Whitney as not normally distributed).

We next sought to identify whether overexpression of Myc and Cyclin T1 with M-C modRNA could regenerate the adult murine heart after MI. Adult CD1 mice were infarcted and intramyocardially injected with 100 µg of M-C, FLuc or GFP modRNA during surgery, immediately post-MI. M-C modRNA injected mice demonstrate greater cardiac function at 28 days post MI compared to control (Figure 6C and D). The change in LVEF (ΔEF%) at day 28 post-MI for FLuc and M-C were −37±3 and −26±3 %, respectively (Figure 6D, Supplemental Figure 6B). At day 28 post MI there was an increase in relative heart weight (Figure 6E), no change in cardiomyocyte cross sectional area (Figure 6F) and scar size (Figure 6G and H). However, we observed an increased left ventricular muscle area (Figure 6I) at 28 days post-MI in M-C modRNA injected mice suggesting that acute M-C modRNA treatment immediately post-MI may induce a partial regenerative response. The increase in left ventricular size in the absence of alterations in cardiomyocyte cross-sectional area prompted us to establish if M-C modRNA treatment provided cardioprotective effect or drives cardiomyocytes into cell cycle. A cohort of mice were infarcted, intramyocardially injected with GFP or M-C modRNA and collected at 4 hours post MI. We observed no change in apoptosis determined by Cleaved caspase 3 and TUNEL staining between the cohorts indicating M-C does not drive a protective effect (Figure 7A). Additional cohorts of mice were infarcted, intramyocardially injected with FLuc or M-C modRNA and pulsed with EdU at 8 and 24 hours (Figure 7B). M-C modRNA significantly increased cardiomyocyte DNA synthesis (EdU) and mitosis (P-H3) (Figure 7C and D). Furthermore, at day 5 post-MI total heart cardiomyocyte number was increased (Figure 7E) indicating that M-C modRNA drives proficient cell division and cardiac regeneration, similar to observations in MycER^T2^ + Cyclin T1 expressing mice.

**Figure 7:**
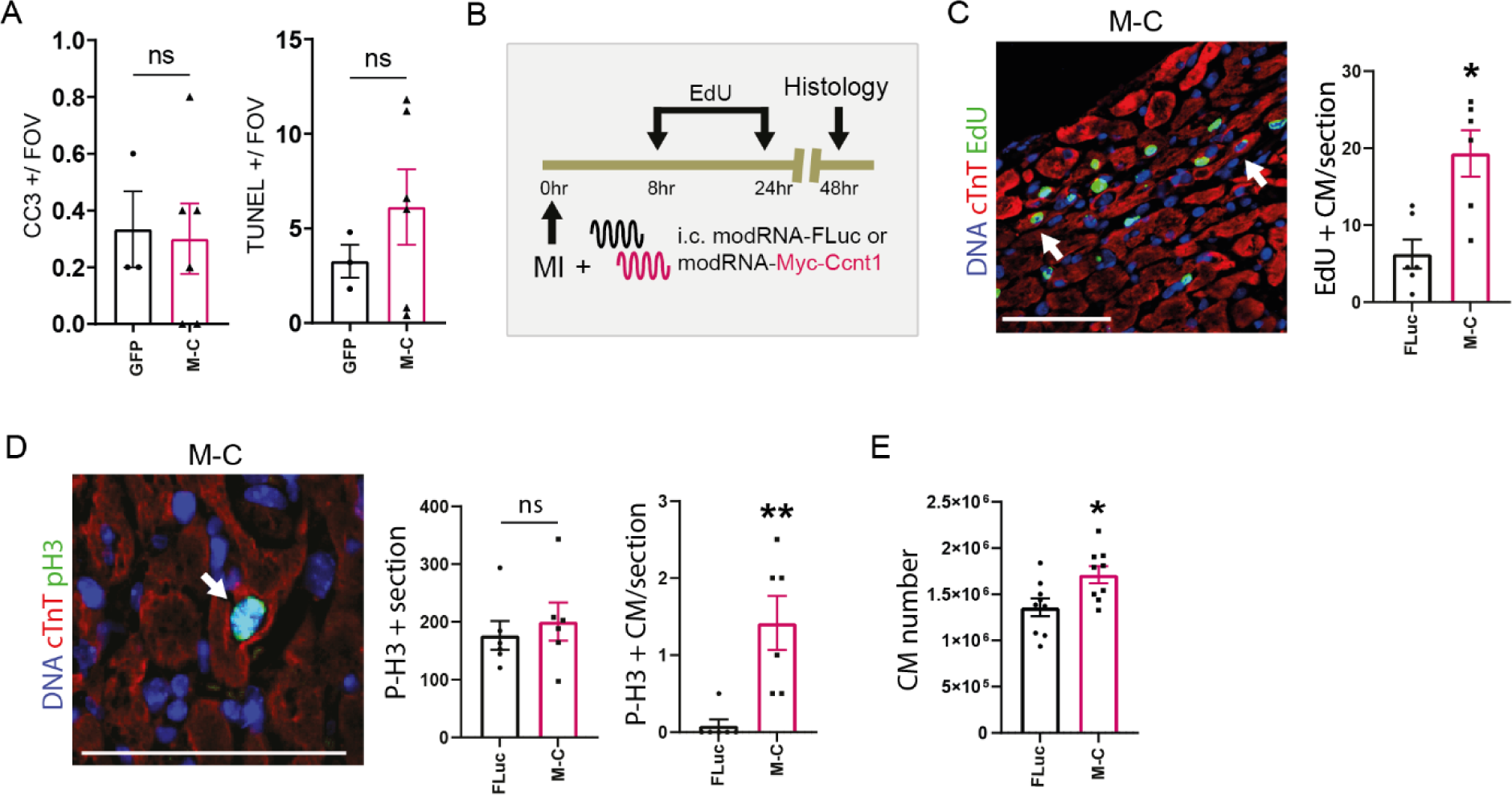
Assessment of cardiomyocyte proliferation following MI and M-C injection in mice. (A) Quantification of cleaved caspase 3 and TUNEL positivity from GFP (n = 3) and M-C (n = 4) injected mice at 4-hour post modRNA injection. (ns-not significant) (B) Experimental schematic to assess M-C induced cardiomyocyte proliferation. At the time of MI, each mouse was injected with 100 µg of FLuc or M-C. Then each mouse was intraperitoneally injected with 12.5 mg/kg EdU at 8- and 24- hours post MI and collected at 48 hours post MI. (C) Immunofluorescence staining and quantification of EdU (green), cardiac troponin T (Red) and DNA (Blue) in FLuc and M-C injected hearts following administration of EdU at 8 and 24 hours. p = 0.011, n = 6. (D) Immunofluorescence staining and quantification of mitotic cardiomyocytes (P-H3+/Tnnt2+) of P-H3 (green), cardiac troponin T (Red) and DNA (Blue) in M-C and FLuc injected heart at 48 hours post modRNA injection. Mitotic cardiomyocytes were increased, p = 0.007, n = 6, numbers of non-myocyte P-H3+ were unchanged (ns- not significant) (E) Quantification of cardiomyocyte number of M-C and FLuc injected hearts 5 days post MI. Unpaired t-test = 0. 0.0176, n = 9. Mann-Whitney tests were performed for all statistical comparisons unless stated due to low n. Scale bars are 100µm.

## Discussion

Heart failure is a clinically unmet need with no widely adoptable therapy, therefore, developing therapeutics to re-muscularize the heart is of the utmost importance. It is now widely accepted that in regenerative species, or regenerative developmental stages such as the neonatal mouse, the majority of regenerated cardiomyocytes in the heart descend from pre-existing resident cardiomyocytes and are derived by cardiomyocyte proliferation^16, 17^. These studies also establish that forced cardiomyocyte proliferation is a viable strategy for myocardial regeneration. However, whether activation of cardiomyocyte proliferation is an effective and safe approach to regenerate the heart and how best to drive regenerative programmes is largely unknown. Towards this goal, the utility of expressing sustained either microRNA-199a or Sav–short hairpin RNA from AAV vectors to stimulate cardiac repair by cardiomyocyte proliferation was recently shown in pigs^18–20^, however, the long-term expression of microRNA-199a eventually resulted in arrhythmic death. The success of endogenous proliferative techniques will likely depend on several factors including genetic strategy, magnitude of response, treatment duration, treatment localisation and treatment timing.

Here we use a genetically modified mouse that carries a cardiomyocyte-specific, switchable version of Myc combined with an AAV that drives cardiomyocyte-specific expression of Cyclin T1 to assess whether dual expression of Myc-Cyclin T1 can be utilised for therapeutic benefit post-cardiac injury. Using a combination of histological techniques and snRNAseq we demonstrate that ectopic expression of Myc and Cyclin T1 drive cardiomyocyte cell-cycle transcriptional programmes that lead to cardiomyocyte proliferation *in vivo*. Combined transient Myc and Cyclin T1 expression with permanent LAD coronary artery ligation resulted in similar levels of Ki67 (6.2%) and P-H3 (2.9%) staining in the RZ to that observed in the undamaged heart^4^. However, the proliferative response was significantly increased in the region adjacent to the infarct (Ki67, 25.1% and P-H3, 6.9%) indicating the BZ provides additional mitogenic stimuli that can increase the ability of MycER^T2^ to drive cardiomyocytes into cycle following MI. We have shown that transiently activating MycER^T2^ combined with Cyclin T1 expression in an undamaged heart drives a proliferative response that does not cause lethality^3^. Unfortunately, combining whole heart transient MycER^T2^ and Cyclin T1 expression with MI caused rapid lethality. We were unable to assess the physiological reason for the death as the mice deteriorated rapidly, no cardiac rupture was observed. Sudden cardiac death may be due to the increased levels of proliferating cardiomyocytes which must undergo sarcomere disassembly before cytokinesis, which we and others have observed in genetic models with enhanced cell-cycle activity^3,4,21–23^. Modelling of cardiomyocyte cell division in hiPSC-CMs has shown that sarcomere breakdown is predominantly observed from pro-metaphase to telophase and that cardiomyocytes briefly stop contracting during cytokinesis, after which sarcomeres are restored and beating is reinitiated ^24^. *In vivo*, cardiomyocytes are electrically and mechanically interconnected, and therefore any interruption of beating is likely to disrupt cardiac output or be an arrhythmogenic risk. The magnitude and location of proliferation initiated within a regenerating heart is likely to be a key factor to determine the most efficacious regenerative therapy while minimising its side effects.

To achieve a pattern of high-level localised and transient MycER^T2^ expression from the genetically encoded *Gt(ROSA)26Sor^tm3(CAG-MYC/ERT2)Gev^* allele, we employed Cre modRNA cardiac injection with tamoxifen treatment to activate MycER^T2^ for 48 hours. Using a genetic reporter, we determined that cardiac injection of Cre allows recombination of ∼9% of the right ventricle and that GFP and FLuc encoding modRNAs drive rapid, transient, and localised expression of proteins. When we combined ectopic cardiomyocyte Cyclin T1 expression with localized, transient Myc expression, lethality was prevented and transient Myc expression led to a 14% increase in ΔEF and a 6.5% reduction in the scar, compared to controls, suggesting that localised, transient, expression post-MI can induce a regenerative response.

Towards clinical translation, we designed a modRNA as a prototypical therapeutic agent for combined Myc-Cyclin T1 delivery to the experimentally infarcted mouse heart. ModRNAs allow the ectopic delivery of genes into cells. Once inside a cell the modRNAs are encoded into the protein of interest by the endogenous translation machinery. The nucleotide modifications allow a change in RNA secondary structure that prevents recognition by the innate immune system and RNA degradation. It is well established that modRNAs are taken up effectively by cardiac tissue and drive transient and robust expression of proteins^13, 25, 26^. Crucially, modRNA delivery is transient, naturally limiting the duration of treatment and allowing for short-term, localised, intense expression of protein. In addition, modRNA has no gene packaging capacity and induces minimal immunogenic response compared to viral vectors which have the potential for random genome integration and ineffective translation due to the presence of pre-existing neutralising antibodies. The clinical potential of modRNAs has recently been demonstrated by the success of modRNA-based SARS-CoV-2 vaccines^27, 28^ and modRNA therapeutics are now an established therapeutic modality. Furthermore, in cardiovascular disease, phase II clinical trials using VEGF-A modRNA to improve vascularisation have met the primary endpoint of safety and tolerability in heart failure patients (NCT03370887)^13, 29^. We show, M-C modRNA drives transient expression of Myc and Cyclin T1 protein and downstream Myc programmes in hESC-CMs *in vitro.* In mice, following an experimental MI and injection of M-C modRNA, we observe Myc and Cyclin T1 protein expression at 1 and 4 hours at the injection site, which is absent by 24 hours. Cardiomyocytes are driven into the cell cycle, leading to increased cardiomyocyte cell number, and improved left ventricular function. These findings are remarkable given that, even with a single dose of transient expression, the regenerative therapeutic initiates the Myc downstream signalling cascade of cellular signalling that leads to cell division.

We observe a short expression period of Myc protein following modRNA transfection (*in vitro* and *in vivo*) which is diminished by 24 hours and extends upon proteasome inhibition. In an endogenous setting, the half-life of Myc is estimated to be around 20 to 40 mins^14, 15^, and Myc is then targeted for degradation by the ubiquitin-proteasome pathway, via the E3 ubiquitin ligase FBXW7^30^. Here we used the endogenous mouse sequence of Myc to design our modRNA and the results show the level and duration of expression rely on the endogenous mechanisms that control the turnover of the protein – which is characteristically rapid for Myc. It is reasonable to hypothesise that modRNA therapeutic designs could be optimised to exploit approaches to stabilise or limit the half-life of ectopically expressed modRNA therapeutic proteins. For cardiac regeneration, both the pulse-like characteristic of modRNAs and the expression of endogenously short-lived proteins, such as Myc, may prove beneficial for cardioregenerative therapies that have a deleterious effect on cardiac function following chronic expression^3,31^. Rapid kinetics is also beneficial from an off-target safety perspective, as modRNAs driving mitogenic programmes would only induce limited round(s) of cell-cycle activity. Furthermore, using unbiased approaches of single nuclei sequencing and proteomics, we demonstrate that a short pulse of ectopic Myc and Cyclin T1 expression drives cardiomyocyte cell-cycle transcriptional programmes that lead to cardiomyocyte proliferation and highlight key processes activated by Myc-Cyclin T1, including the coupling of mitochondrial respiration and glucose metabolism and activation of mTORC1 signalling which may be key to cell cycle entry induced by Myc.

Here, we directly inject unformulated modRNA into mouse hearts at the time of MI. Direct injection of modRNA allows for localisation of RNA to the cardiac tissue and we observed only infrequent recombination events in other organs. This method of application has been used in cardiovascular disease previously (VEGF-A) and direct injection allows for localised transient heart-specific expression and demonstrates the safety and feasibility of the modRNA approach in heart disease^25, 29^. Conceivably, increased efficacy could be achieved through repeated dosing of M-C modRNA to drive further regeneration in the heart. Catheter delivery could be modified and used to achieve local delivery to the heart muscle. Alternatively, further refinement of modRNA delivery platforms may be required to achieve cardiac targeting following systemic delivery.

Myc-Cyclin T1 modRNA carries great potential in addressing the gap in the current therapies available to clinicians. The clinical translation of our therapeutic will depend on answers from further pre-clinical models that will determine the arrhythmic safety profile, determination of off-target toxicity and optimal dosages and timing to achieve meaningful recovery. Together our results demonstrate that localized and transient delivery of cardiac regenerative genes via modRNA is an important therapeutic strategy for cardiac regeneration and that a Myc-Cyclin T1 co-expression transcript has the potential to be an effective medicament via this approach.

## Methods

### Ethical approval and animal experiments

Human tissue experiments were performed with QIMR Berghofer Medical Research Institute (approval number P2385). Animal experiments received ethical approval and were conducted in accordance with either The University of Queensland Animal Ethics Committee (approval number # 2019/AE000149) or The Home Office UK guidelines, under project licence PP2054013 (C.H.W) that were evaluated and approved by the Animal Welfare and Ethical Review Body at the University of Cambridge. All animals were kept under SPF conditions. Mice were maintained on regular diet in a pathogen-free facility on a 12-hr light/dark cycle with continuous access to food and water. All mice were euthanised by a schedule 1 method. Mouse strains *Tg(Myh6-cre)1Jmk/J* and *Gt(ROSA)26Sor^tm14(CAG-tdTomato)Hze^* were obtained from the Jackson Laboratory. *Gt(ROSA)26Sor^tm3(CAG-MYC/ERT2)Gev^* was generated previously^3^. CD-1 mice were obtained from Charles River or Animal Resources Centre. Ear biopsies were collected from 2–3-week-old mice and genotyped by PCR with the following oligonucleotide primers: for *Rosa26CAG*; Universal forward: 5’-CTCTGCTGCCTCCTGGCTTCT-3’ Wild-type reverse: 5’-CGAGGCGGATCACAAGCAATA-3’ and CAG reverse: 5’ TCAATGGGCGGGGGTCGTT-3’. Generic primers to recognise Cre were Cre1: 5′-GCTGTTTCACTGGTTATGCGG −3′ and Cre2: 5′-TTGCCCCTGTTTCACTATCCAG - 3′. Primers to recognise Myc-ER and Tomato were: MycER^T2^-Forward 5’- ATTTCTGAAGACTTGTTGCGGAAA-3’ and MycER^T2^-Reverse 5’- GCTGTTCTTAGAGCGTTTGATCATGA-3’; Tomato-Forward: 5’- CAACTGCCCGGCTACTACTA-3’ Tomato-Reverse: 5’- CCATGTTGTTGTCCTCGAG −3’. AAV9-cTnT-3xFLAG-mCCNT1-WPRE was obtained from Vector Biolabs. For systemic infection, AAV viruses were diluted in PBS, each mouse received 1 × 10^11^ GC in 100 µl via tail vein injection at 4–5 weeks of age. To activate MycER^T2^ 1 mg of tamoxifen (Sigma, T5648) was i.p injected into adult mice in 10 % ethanol and corn oil (10 mg/ml) twice over a 48-hour period and tissues collected at 48 hours post initial i.p. injection.

### Adult mouse MI and intracardiac modRNA injections

Mice were anaesthetised with 2% isoflurane (Bayer) and intubated. Ventilation parameters were as follow: 0.4 L/min oxygen, tidal volume of 250 μL and a respiration rate = 133 strokes/min (Minivent, Harvard Apparatus). To maintain body temperature, mice were positioned on a heated surgical mat and secured with surgical tape. The anteriolateral chest was shaved and a lateral thoracotomy of the 4^th^ intercostal space was performed. The pericardium was blunt dissected with fine nosed tweezers. The left anterior descending coronary artery was permanently ligated with 7-0 prolene suture, just below the LAD exit from the underneath the left atria. MI was verified by observing blanching of the myocardium. Transfection mix of modRNAs were composed of 100 µg modRNA in 20 µl PBS and 40 µl RNAiMAX reagent. 20 µl of transfection mix was injected at 3 locations in the ischaemic penumbra; injections in the superior anterior, inferior anterior and superior posterior left ventricular wall. The chest wall was then closed with 4-0 silk suture and skin closed with 6-0 prolene suture. For post operative analgesia, buprenorphine (0.1 mg/kg) and carprofen (10 mg/kg) was injected subcutaneously. For EdU, mice were intraperitoneally injected with 12.5 mg/kg EdU at 8 hours and 24 hours post MI.

### ModRNA

mMyc-T2A-Ccnt1 (Supplemental data), Cre and luciferase modRNA was synthesised by Trilink Biotechnologies. For eGFP and hMYC-T2A-CCNT1 (Supplemental data) modRNA synthesis, 1 µg of template plasmid was digested using 1µL AgeI (NEB, R3552S) for 15h in Cutsmart Buffer (NEB, B7204S) and purified using QIAGEN QIAquick PCR Purification Kit (Qiagen, 28104) according to the manufacturer’s protocol. Template for modRNA synthesis was PCR amplified from linearised plasmid using Platinum Taq DNA Polymerase High Fidelity (Invitrogen, 11304-011) using manufacturer’s programme specifications and the following primers.

Primer sequences (5’-3’): PolyT primer: TTTTTTTTTTTTTTTTTTTTTTTTTTTTTTTTTTTTTTTTTTTTTTTTTTTTTTTTTTTTTTTTTTTTTTTTTTTTTTTTTTTTTTTTTTTTTTTTTTTTTTTTTTTTTTTTTTTTTTTTTCCCTCACTTCCTACTCAGGC, Fwd T7 primer: GTGAGCGCGCGTAATACG PCR product was purified using QIAquick PCR Purification Kit. modRNA was generated by in vitro transcription using MEGAscript T7 Transcription Kit (Invitrogen, AM1333) with 6 to 24h incubation time. Reaction was set up using ARCA (NEB, S1411L or Trilink, N-7003-5) and N1-Methylpseudouridine-5’-Triphosphate (Trilink, N-1081-5) to replace UTP. After incubation, synthesis products were treated with TURBO DNase for 15 min and purified using MEGAclear Transcription Clean-Up Kit (Invitrogen, AM1908) according to the manufacturer’s instructions. Subsequently, modRNA was treated with Antarctic Phosphatase (NEB, M0289S) for 1h. It was then re-purified with the MEGAclear kit, and the concentration was quantified using a NanoDrop spectrometer (Thermo Scientific). To be used in intramyocardial injections, modRNA was concentrated using Amicon Ultra-0.5 Centrifugal Filter Unit 30kDa (MilliporeSigma, UFC5030) at 4°C at 12 000g for 10 min to yield 100 µg in 20µl and combined with 40 µl Lipofectamine RNAiMAX Transfection Reagent (Life Technologies, 13778075).

### Echocardiography

Cardiac function was assessed on day 0, 3, 6 or 14 and 28 post MI surgery. Researchers were blinded to genotypes/treatment. Echocardiographical measurements were obtained using a Vevo 2100 or 3100 ultrasound system fitted with a MS550D transducer, 40 MHz center frequency (Fujifilm Visualsonics). Mice were anaesthetised with isoflurane (1.5% at 1 L oxygen / min). Body temperature was maintained with a heated stage during echocardioagraphy. Heart and respiration rates were monitored with ECG pads. Cardiac function was estimated either using B-Mode images of the left ventricle, which were obtained from three short-axis views (proximal, mid and distal positions) and one parasternal long-axis view, or M-Mode images of the parasternal long-axis and short-axis views tracing measurements over 3 to 6 contraction periods. For each view, the endocardial surface was traced and measurements in systole and diastole were used to calculate ejection fraction in Vevolab analysis software v3.2.6 (Fujifilm Visualsonics).

### Bioluminescence imaging

Luciferase-mediated bioluminescence was measured at 8h, 24h, 48h and 72h using an IVIS Spectrum (Perkin Elmer, USA). Mice were injected intraperitoneally with D-luciferin diluted in PBS (15 mg/ml stock) at 150 mg/kg, anaesthetised and imaged until the bioluminescence plateaued. Typical settings were medium binning, f-stop 1, exposure time 60 seconds, field of view (FOV) 13cm, and open field. Average radiance (photons/second/cm2/sr) of regions of interest (ROI) drawn over the chest was determined using Living Imagine software v4.5.4 (Perkin Elmer, USA).

### Isolation of adult cardiomyocytes

Fix-dissociation of cardiomyocytes. Whole hearts were isolated from mice and washed in PBS followed by fixation in 1% PFA overnight at room temperature with agitation. The following day hearts were washed four times in PBS. Hearts were cut into 1–2 mm^3^ pieces and incubated with 0.5 U/mL collagenase B (Roche #11088807001) in 0.2% NaN_3_/PBS and left to oscillate at 1000 rpm at 37 °C. Every 12 h, the cardiomyocyte supernatant was collected and stored at 4 °C in 0.2% NaN_3_/FBS. Once dissociation was complete (∼8 days) cells were centrifuged at 1000 × g for 3 min, washed twice in PBS and resuspended in 0.2% NaN_3_/PBS at 4 °C. Cardiomyocytes were then counted using a haemocytometer.

### hESC-CM generation

Ethical approval for the use of human embryonic stem cells (hESCs) was obtained from QIMR Berghofer’s Ethics Committee (P2385) and was carried out in accordance with the National Health and Medical Research Council of Australia (NHMRC) regulations. Cardiomyocyte/stromal cell cultures were produced from HES3 human embryonic stem cells (hESCs, WiCell) maintained using mTeSR-1 (Stem Cell Technologies). hESCs were seeded at 2×10^4^ cells/cm^2^ in Matrigel coated flasks and cultured for 4 days using mTeSR-1 and passaged with TrypLE (ThermoFisher). Subsequently, hESCs were differentiated into cardiac mesoderm by culturing for 3 days in RPMI B27-medium (RPMI 1640 GlutaMAX + 2% B27 supplement minus insulin (ThermoFisher), 200 μM L-ascorbic acid (Sigma), 1% Penicillin/Streptomycin (ThermoFisher)) and growth factors: 5 ng/ml BMP-4 (R&D Systems), 9 ng/ml Activin A (R&D Systems), 5 ng/ml FGF-2 (R&D Systems) and 1 μM CHIR99021 (STEMCELL Technologies). Media was exchanged daily. Cardiomyocyte specification was completed by 3 days of culture in RPMI B27-media with 5 μM IWP-4 (STEMCELL Technologies), followed by another 7 days of culture with RPMI B27+ (RPMI 1640 GlutaMAX + 2% B27 supplement with insulin (ThermoFisher), 200 μM L-ascorbic acid (Sigma) and 1% Penicillin/Streptomycin (ThermoFisher)). The cardiac cells, comprised of ∼70% cardiomyocytes and, ∼30% CD90+ stromal cells were dissociated at 15 days using human cardiac digestion buffer: 0.2% collagenase type I (Sigma) in 20% fetal bovine serum (FBS) in PBS (with Ca2+ and Mg2+) for 45 min at 37°C. The cardiac cells were further digested with 0.25% trypsin-EDTA for 10 min. The cells were strained through a 100 µm filter, centrifuged at 300 x g for 3 min, and resuspended at the required density for 3D culture in CTRL Medium: α-MEM GlutaMAX (ThermoFisher), 10% FBS, 200 μM L-ascorbic acid (Sigma) and 1% Penicillin/Streptomycin (ThermoFisher).

H9/WA09 hESCs, were maintained in TeSR-E8 (Stem Cell Technologies) at 37°C. Media was changed daily and cells were passaged using ReLeSR (Stem Cell Technologies). For differentiation, hESCs were dissociated using TrypLE (ThermoFisher) and seeded at 1 million or 500,000 cells/well in Matrigel coated 6 well or 12-well plates, respectively. hESCs were differentiated using a protocol previously described^32^. Briefly, mesoderm was induced by CDM-BSA media supplemented with FGF2 (20 ng/ml, Qkine), Activin-A (50 ng/ml, Qkine), BMP4 (10 ng/ml, Bio-Techne) and Ly294002 (10 ng/ml, Alfa Aesar) for 42 hours. Media was changed to CDM-BSA supplemented with FGF2 (8 ng/ml), BMP4 (10 ng/ml), retinoic acid (1 µM, Sigma Aldrich), and IWR1 (1 µM, R&D Systems) for 48 hours and was refreshed for a further 48 hours. Then, media was changed to CDM-BSA supplemented with FGF2 (8 ng/ml) and BMP4 (10 ng/ml) for 48 hours, then CDM-BSA without any supplements was refreshed every 48 hours to maintain the cardiomyocytes. After beating was observed, lactate selection was carried out as previously described (usually on day 14 or day 15 since starting differentiation). Briefly, cardiomyocytes were replated at 2 million or 1 million cells/well in Matrigel coated 6 well or 12 well plates. The following day, media was changed to DMEM without glucose/pyruvate supplemented with sodium lactate (4 mM, Sigma Aldrich) for 48 hours and refreshed for a further 48 hours. Cardiomyocyte purity was within the range of 70-95% measured by Troponin T flow cytometry.

### hESC-CM 2D cell culture

Day 15 dissociated differentiation cultures were plated at 150,000 cells/cm^2^ or 100,000 cells/cm^2^ in 0.1% gelatin-coated 96-well or 12-well tissue culture plates in CTRL medium, respectively, and cultured for 2 days. Cardiac cells were then cultured in Maturation Media (DMEM, no glucose, no glutamine, no phenol red (ThermoFisher), 1% P/S (ThermoFisher), 200 mM ASC (Sigma), 1% GlutaMAX (ThermoFisher), 4% B27-(ThermoFisher), 100 µM Palmitate (Sigma), 1 mM Glucose (Sigma), 33 ng/mL Aprotinin (Sigma)) for 3 days.

For transfection, 1 µg or 4 µg of modRNA was mixed with Opti-MEM reduced serum medium, GlutaMAX supplement (ThermoFisher) and Lipofectamine RNAiMAX (Invitrogen) as per manufacturer’s instructions for 96-well or 12-well tissue culture plates, respectively. modRNA/lipid complex was added to cells 1:10 with Maturation Media and gently rocked to ensure even spreading of transfection complex. For proliferation experiments, cells were cultured for 48 hours whilst for qPCR and Western Blotting, cells were cultured for times indicated.

### Mouse embryo fibroblasts

MEFs were generated from embryos 13.5 d.p.f. and cultured in DMEM (Thermo Fisher 41966052) supplemented with L-glutamine (Thermo Fisher, 25030-024) and 10% BGS (Hyclone SH30541.03HI). For experiments, MEFs were plated onto coverslips at 1.5×10^4^ cells per well of a 24-well plate in 10% BGS lacking Tamoxifen, of which they were starved for three days to render cells quiescent. After 72h of Tamoxifen withdrawal, MEFs were transfected with Lipofectamine RNAiMAX Transfection Reagent according to the manufacturer’s instructions for delivery of 1 µg (eGFP) or 5 µg of modRNA (M-C human or mouse). Mock transfections lacking RNA were performed as negative controls.

### Immunostaining of 2D cultures

Cardiac cells were fixed with 1% paraformaldehyde (Sigma) for 20 min at room temperature and washed 2x with PBS after which they were incubated with Blocking Buffer, 5% FBS and 0.2% Triton X-100 (Sigma) in PBS, for 45 min at room temperature. Cells were incubated with mouse anti-α-actinin (Sigma, A7732, 1:1000) and rabbit anti-Ki67 (Cell Signaling, #9129S, 1:400) diluted in Blocking Buffer for 1-2 hours at room temperature. Cells were then washed 2x with Blocking Buffer and subsequently incubated with Alexa Fluor 633 goat anti-mouse IgG (Invitrogen, A21050, 1:400), Alexa Fluor 555 goat anti-rabbit IgG (Invitrogen, A21429, 1:400) and Hoechst33342 (Invitrogen, H1399, 1:1000) diluted in Blocking Buffer for 1-1.5 hours at room temperature. They were then washed 2x with Blocking Buffer and imaged. For screening, cardiac cells were imaged using Nikon ANDOR WD Revolution Spinning Disk confocal microscope. Custom batch processing files were written in MATLAB R2013a (Mathworks) to analyze cardiomyocyte number, identify nuclei of proliferating (Ki67^+^) cardiomyocytes and determine average cardiomyocyte size, and export batch data to an Excel (Microsoft) spreadsheet.

For BrdU pulse experiments, cells were fixed in cold MetOH 24h pos-transfection with modRNA and stored at −20°C until processed for immunostaining. Specifically, coverslips were rehydrated with PBS for 10 minutes, followed by antigen retrieval by denaturing DNA with 2N HCl, 0.2% Triton X for 15 minutes at 37°C. This was followed by 3x washes in PBST (PBS with 0.1% Tween), incubation in 50mM glycine in PBS for 5 minutes, and 3 more PBST washes. Coverslips were incubated in blocking solution (2.5% goat serum, 1% BSA in PBST) for 20 minutes. Overnight incubation with primary antibody against BrdU (Rat monoclonal [BU1/75(ICR1)]; Abcam ab6326; 1:200) diluted in blocking was carried out at 4°C and followed by 3x washes in PBST. Secondary antibody (Goat anti-Rat IgG (H+L) Cross-Adsorbed Secondary Antibody, Alexa Fluor 488) was diluted in blocking solution and incubated for 1h at room temperature. Coverslips were washed 3x 5 minutes in PBST, with the second wash containing Hoechst (1:5000) and mounted on slides using ProLong Gold Antifade Mountant (Thermo Fisher, P36934). Image analysis was performed on Cell Profiler where Hoechst- and BrdU-positive nuclei were segmented and counted using the “Identify Primary Objects” function. Specifically, global Minimum Cross-Entropy thresholding was applied. For CMs, where segmentation wasn’t efficient in regions with CM clusters, manual counting of the positive nuclei in the clusters was performed.

### Nuclei isolation method for snRNAseq

Cardiac nuclei isolation method was adapted with modifications from literature ^33, 34^. Whole heart was dissected into 1mm pieces on ice with scalpel in 8ml of buffer (300 mM sucrose, 5mM CaCl2, 5 mM Mg acetate, 2mM EDTA, 0,5mM EGTA, 10 mM Tris-HCl, 1 mM dithiothreitol (DTT), 1× protease inhibitor, 0.4 U μl−1 RNaseIn in nuclease-free water) and transferred into the dounce, homogenised over 10 strokes with pestle A and 8 with pestle B on ice, strained through 40um cell strainer and centrifuged 500g 5min at 4C. Further supernatant was discarded, and pallet was resuspended in 0,5ml 1M sucrose buffer (1 M sucrose, 5 mM Mg acetate, 10 mM Tris-HCl, 1 mM dithiothreitol (DTT), 1× protease inhibitor, 0.4 U μl−1 RNaseIn in nuclease-free water) and layered over 1ml 1M sucrose buffer, centrifuged at 1000g 5min at 4C and resuspended in 0,5ml PBS+ 2%BSA + SUPERaseIn with 1:1000 Hoechst. FACS sorting of nuclei was performed using the SH800Z cell sorter (Sony), after which nuclei were collected by centrifuging 600g 8min 4C, counted using hemocytometer and dilute to yield 10 000 nuclei in 43ul 1x PBS, 2%BSA, 0.2 U μl−1 SUPERaseIn.

Nuclei were submitted to the Cancer Research UK (CRUK) Cambridge Institute for 10x library preparation using V3 chemistry and sequencing on a NovaSeq 6000 (Illumina).

### snRNAseq processing and data analysis

Alignment was performed using Cellranger mkfastq to mouse mm10/GRCm38 pre-mRNA reference transcriptome to generate transcript count matrix. Processing and analysis done in R, using Seurat v4.0.0. package. Low-quality cell filtering was based on 500<genes<2500, mt.percentage<10%. dimensional reduction performed according to Seurat sctransform pipeline, with resolution parameter assessed using clustree package. Cluster identification was done using FindClusters and manually annotated using marker genes^35, 36^. Doublet filtering performed using DoubletFinder, assuming 5-20% doublets, according to predicted recovered nuclei number. Data integration done using RPCA algorithm, differential gene expression (DGE) analysis done by Seurat FindMarkers (Wilcox test, log2FC > 0.25 positive markers with p_adj < 0.01) on RNA assay normalised and scaled counts. Gene set enrichment analysis done using Broad Institute run MSigDB database. For analysis of cardiomyocyte signatures, cardiomyocyte clusters from all datasets were subset and reintegrated, before proceeding with subclustering and marker identification. We performed Seurat CellCycleScoring and identified pseudotime cell trajectory with Monocle 3.

### Time course proteomics

2D cardiomyocytes grown in 6-well plates were transfected with Lipofectamine RNAiMax with 10 µg MYC-CCNT1 (M-C) modRNA or without modRNA (CTRL). At 1, 4, 18 and 48 hr timepoints, cells were lysed in RIPA Buffer (150 mM NaCl, 5 mM EDTA, 1% Triton X-100, 0.5% Sodium Deoxycholate, 0.1% SDS and 50 mM Tris, pH 8.0) supplemented with protease inhibitor cocktail (Roche), scraped into 1.5 mL Eppendorf tubes. Proteins were quantified by Pierce BCA Protein Assay kit (Thermo Scientific), and 35 µg were aliquoted for proteomics. No significant difference was found in protein concentration between M-C and CTRL conditions at any time point (Figure 5E). Proteomics samples were reduced with 5 mM dithiothreitol (Sigma-Aldrich) at 60° C for 30 min and alkylated with 10 mM Iodoacetamide (Sigma-Aldrich) for 10 min at room temperature in the dark. Proteins were precipitated with 5x ice cold acetone (HPLC grade, Sigma-Aldrich) and centrifuged at 20,627 g for 10 min at 4° C. The supernatants were removed, and the pellets were washed three times with acetone. Samples were then reconstituted in 40 mM Triethylammonium bicarbonate buffer (TEAB, Sigma-Aldrich) and digested overnight with Porcine trypsin (Promega) at 37° C. Digestion was quenched with trifluouroacetic acid (TFA, Thermo Scientific) and samples were cleaned using self-fabricated Empore SDB-RPS StageTips, dried by GeneVac sample concentrator, then reconstituted in 0.1% formic acid (FA, Fisher Scientific) for LC-MS analysis. Peptide concentration was determined by Pierce MicroBCA Assay kit, and all samples adjusted to 500 ng/µL.

Samples (2 µL injection) were loaded on to a Thermo Acclaim PepMap 100 trap column (5 mm x 300 µm ID) for 5 min at a flow rate of 10 ul/min with 95% Solvent A (0.1% FA in water) and subsequently separated on a Thermo PepMap100 analytical column (150 mm x 300 µm ID) equipped on a Thermo Ultimate 3000 LC interfaced with a Thermo Exactive HF-X mass spectrometer. Peptides were resolved using a linear gradient of 5% solvent B (0.1% FA in 80% acetonitrile) to 40% solvent B over 120 min at a flow rate of 1.5 µL/min. This was followed by column washing and equilibration for a total run time of 65 min. Mass spectrometry data was acquired in positive ion mode. Precursor spectra (350-1400 m/z) was acquired on orbitrap at a resolution of 60,000. The AGC target was set to 3×10^6^ with a maximum injection time of 30 ms. Top 20 precursors were selected for fragmentation in each cycle and fragment spectra was acquired in orbitrap at a resolution of 15,000 with stepped collision energies of 28, 30 and 32. The AGC target was 1e^5^, with a maximum ion injection time of 45 ms. The isolation window was set to 1.2 m/z. Dynamic exclusion was set to 30 sec and precursors with charge states from 2-7 were accepted for fragmentation.

Raw LC-MS data was searched against the reviewed Uniprot human database (20,399 sequences, downloaded April 19, 2021) using Sequest HT on the Thermo Proteome Discoverer software (Version 2.3), with matching between runs enabled. Precursor and fragment mass tolerance were set to 20 ppm and 0.05 Da respectively. A maximum of two missed cleavages were allowed. A strict false discovery rate (FDR) of 1% was used to filter peptide spectrum matches (PSMs) and was calculated using a decoy search. Carbamidomethylation of cysteines was set as a fixed modification, while oxidation of methionine and N-terminal acetylation were set as dynamic modifications. Protein abundance was based on intensity of the parent ions and data was normalized based on total peptide amount. Statistical significance was calculated using a t-test for summed abundance-based ratios, built into the Proteome Discoverer software. Graphics were generated using: GraphPad Prism (version 8.4.3) for proteins of interest, with two-way ANOVA statistical analysis; pheatmap (v.1.0.12) package for R (v.4.0.2) including clustering using Pearson correlation; stats package (v.4.0.2) for PCA analysis; graphs collated using Inkscape software. Functional enrichments utilised the EnrichR web tool for Gene ontology biological process 2021, and MSigDB Hallmark 2020 dataset enrichments.

### RNA isolation

RNA was isolated from 2D cells using RNeasy Micro Kit (QIAGEN) as per manufacturer’s instructions. RNA was isolated from mouse tissues using TRIzol^®^ (ThermoFisher Scientific) as per manufacturer’s instructions. For whole tissues, tissues were homogenised in 1 mL of TRIzol and left at room temperature for 3 minutes to fully dissociate nucleoprotein complexes. 200 µL of chloroform was added, samples were vortexed for 30 seconds, incubated for 3 minutes at room temperature and centrifuged at 12,000 x g for 15 minutes at 4°C. The aqueous phase containing RNA was transferred to a new Eppendorf tube, 1 mL of isopropanol was added and 1 µL of GlycoBlue™ (ThermoFisher) was used as a precipitant. The RNA pellet was washed in 75% ethanol/ 25% nuclease free water, allowed to dry and then resuspended in 87.5 µL nuclease-free water and incubated at 55°C for 15 minutes. RNA was quantified using NanoDrop 2000 spectrophotometer (ThermoFisher Scientific).

### cDNA generation and qPCR

For cDNA generation, RNA was first treated with RNase-free DNase (QIAGEN) and then column-isolated using RNeasy MinElute Clean-up kit (QIAGEN) as per manufacturer’s instructions. RNA was eluted from MinElute columns with 20 µL of nuclease-free water. RNA was then used for qPCR. Superscript III first-strand synthesis system (Invitrogen) was used to generate cDNA. DNase-treated RNA was pre-hybridised with 1 µL 150 ng random hexamer primers and 1 µL 10 mM dNTPs at 65°C for 5 minutes and placed on ice. The SuperScript^®^ reaction mix was added (4 µL of 5x First strand buffer, 1 µL of 0.1 mM dithiothreitol and 1 µL of RNase OUT (ThermoFisher) and 1 µL of SuperScript^®^ III (ThermoFisher). Reverse transcriptase free controls had all the same reagents but without SuperScript^®^ III. cDNA was generated using a thermocycler at 25 °C for 10 minutes, 50 °C for 60 minutes and 70 °C for 15 minutes.

qPCR was performed using SYBR™ Green PCR Master Mix (ThermoFisher) PowerUp™ SYBR™ Green (ThermoFisher) as per manufacturer’s recommendations. qPCR was performed with a QuantStudio™ 5 system (Applied Biosystems) using manufacturer’s instructions. Endogenous controls for human cells was hypoxanthine-guanine phosphoribosyltransferase (*Hprt or HPRT1*). For mouse tissue endogenous controls were Actin and Gapdh. Primer sequences for qPCR are listed in the Table below.

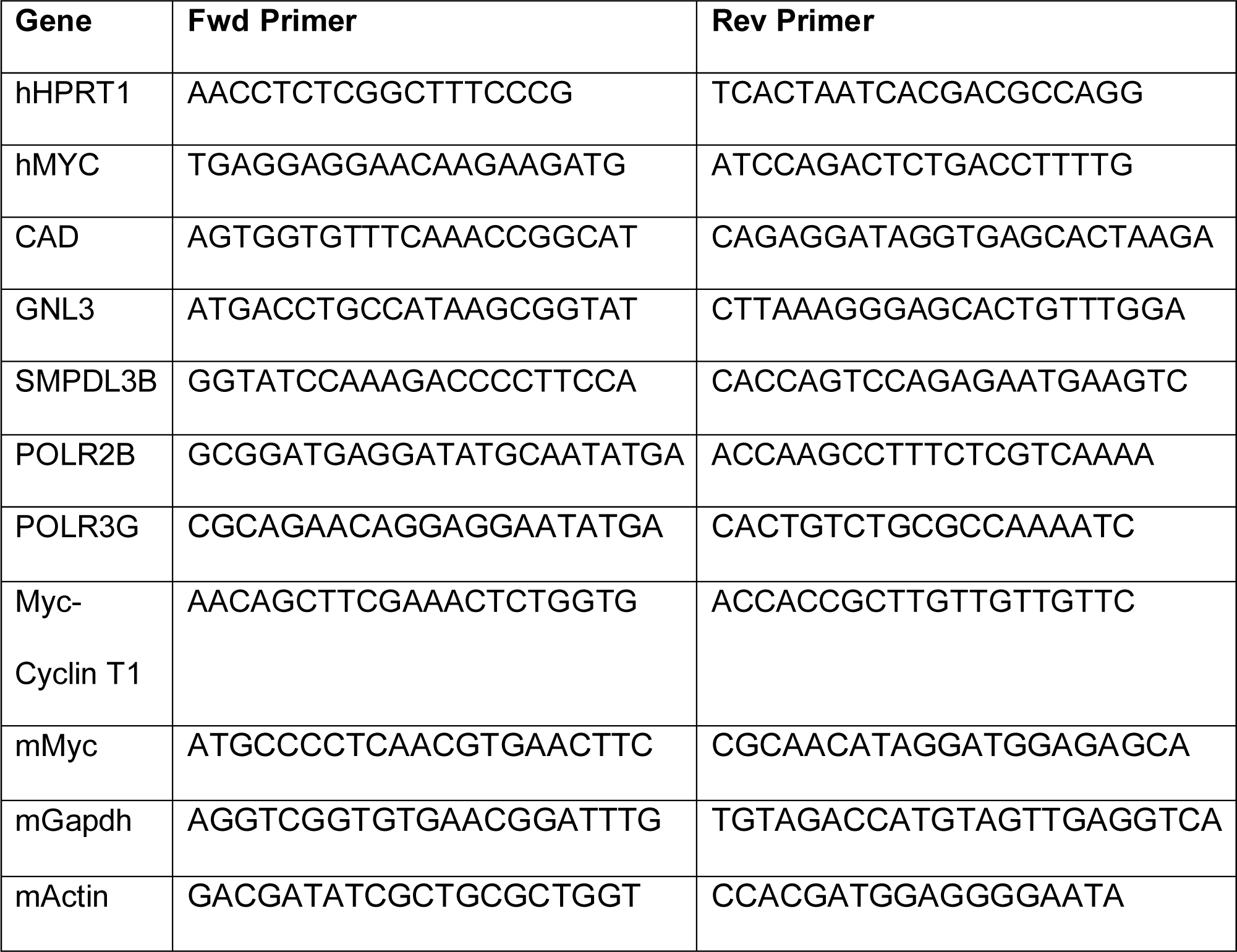

### Western blotting

Cardiac cells were lysed using RIPA Buffer (150 mM NaCl, 5 mM EDTA, 1% Triton X-100, 0.5% Sodium Deoxycholate, 0.1% SDS and 50 mM Tris, pH 8.0) or Lysis buffer (1 % SDS, 50 mM Tris pH6.8 and 10 % Glycerol) supplemented with protease inhibitor cocktail (Roche) and subsequently quantified using Pierce BCA Protein Assay Kit (Thermo Scientific). Samples in lysis buffer were boiled for 10 minutes and sonicated (Bioruptor, Diagenode) for 15 minutes at room temperature, prior to quantification. Following quantification, 6X Loading buffer (0.3 mM Tris pH 6.8, 0.6% Bromophenol blue, 60% Glycerol, 6% 2-mercaptoethanol) with 10% DTT (1M) and/or 10% 2-mercaptoethanol was added and samples were boiled again. Samples in RIPA buffer were denatured by boiling at 70°C for 10 min in 4X LDS Sample Buffer (Invitrogen) and 10X Bolt Reducing Agent (Invitrogen). Proteins were separated by SDS-PAGE using a Bolt 4-12% Bis-Tris acrylamide gel (Invitrogen) or 10% acrylamide/bis gel. Proteins were transferred to a PVDF membrane using Mini Blot Module (Invitrogen). Membranes were blocked in Intercept (PBS) Blocking Buffer (LI-COR) or 5% milk in PBS with 0.1% Tween 20 (PBST), for 1 hour at room temperature then incubated overnight at 4°C with rabbit anti-c-Myc (Abcam, ab32072) or rabbit anti-Cyclin T1 (Cell Signaling, #81464) and mouse anti-GAPDH (Cell Signaling, #97166) diluted 1:1000 in Intercept Blocking Buffer or 5% milk in PBST. After five washes with PBST membranes were incubated with IRDye 680RD goat anti-rabbit IgG (LI-COR, #926-68071) and IRDye 800CW goat anti-mouse IgG (LI-COR, #926-32210) diluted 1:10000 in Intercept Blocking Buffer for 1.5 hour at room temperature or HRP-conjugate antibodies goat anti-rabbit IgG (A0545, Sigma-Aldrich) and anti-mouse IgG (A4416, Sigma-Aldrich). After five washes with PBST and one wash with PBS, fluorescently labelled membranes were dried and imaged using Odyssey CLx Infrared Imaging System (LI-COR); HRP-conjugated membranes were washed three times with PBST and incubated with Immobilon Western Chemilum HRP Substrate (WBKLS0500) and imaged on an Odyssey XF imaging system (LI-COR). Densitometry performed using Image Studio Acquisition Software (LI-COR).

### Histology

Animals were euthanised and hearts were washed in PBS and then fixed either in 10% neutral-buffered formalin or 4% paraformaldehyde/PBS on a shaker overnight at room temperature. The hearts were then washed in PBS, transversely cut with a razor blade and dehydrated in ethanol/xylene and paraffin embedded for sectioning. Tissues were then sectioned at 4 µm and mounted on slides (FisherScientific). For Masson’s Trichrome staining, sections were deparaffinized and rehydrated and stained in Weigert’s iron haematoxylin for 10 minutes and washed in distilled water for 10 minutes. Sections were then stained in Biebrich scarlet-acid fuchsin solution for 10 minutes and further washed in distilled water. Slides were stained in phosphomolybdic-phosphotungstic acid solution for 10 minutes and then aniline blue solution for 0.5-3 minutes (depending on the strength of aniline blue dye). Slides were washed and then differentiated in 1% acetic acid/distilled water solution for 3 minutes. Slides were dehydrated in ethanol/xylene and mounted. For scar and LV muscle quantification, slides 250 µm apart from the aortic valve to the apex were stained with trichrome, imaged on an Aperio slide scanner (Leica). Scar area was calculated as a function of scar area/total heart area. LV muscle area was obtained using ImageJ by drawing around the LV muscle area for each section.

### Immunofluorescent staining of mouse heart sections

For each stain, sections were rehydrated, blocked with 10% goat serum in PBS and stained with primary antibodies in 2% goat serum/PBS overnight at 4°C. Heart sections were washed in PBS stained with secondary antibodies and Hoechst 33342 (Life Technologies) diluted 1:1000 in 2% goat serum/PBS for 1 hour at room temperature and mounted in ProLong Glass mounting media (ThermoFisher). Each slide was then imaged with a Leica Thunder microscope or Zeiss Axio Imager using the Zeiss AxioVision 4.8 software or Leica DM6 B using LASX software. For adult cardiomyocyte proliferation analysis, either 5 images were captured in the border and remote zone and the mean percentage of positive cardiomyocytes was calculated per mouse or 2 entire cross sections for each heart were quantified. For apoptosis analysis, and due to the low level of positivity in the heart, 5 images were captured in the border zone and the mean percentage of positive cells per field was calculated per mouse. Primary antibodies: cardiac troponin T (13-11; Thermo Fisher, MA5-12969; 1:100) and (CT3, Santa Cruz Biotechnology, sc-20025, 1:50). phospho-Histone H3 (Ser10; Merck Millipore, 06-570: Antibody; 1:500), anti-Aurora B antibody (Abcam, ab2254; 1:200), anti-PCM1 (SigmaAldrich, HPA023370, 1:100), anti-Ki67 (SolA15; Thermo Fisher, 1:100) and wheat germ agglutinin, Alexa Fluor™ 488 conjugate or Texas Red-X Conjugate (Thermo Fisher, W11261, W21405). Secondary antibodies were: Goat anti-Rat IgG (H+L) Cross-Adsorbed Secondary Antibody, Alexa Fluor 488 (A-11006); Goat anti-Rabbit IgG (H+L) Cross-Adsorbed Secondary Antibody, Alexa Fluor 488 (A-11008); Goat anti-Rabbit IgG (H+L) Secondary Antibody, Alexa Fluor 647 (A-21244). Hoechst 33258 (bisbenzimide H33258; B2883-100MG; Sigma-Aldrich) was used for DNA stain. TUNEL staining we performed following the manufactures instructions using the ApopTag® Fluorescein In Situ Apoptosis Detection Kit (S7110 SigmaAldrich).

### Immunohistochemistry

Immunohistochemistry was performed on 4 µm sections. Sections were de-paraffinized and rehydrated. Antigens were retrieved by boiling in 10 mM citrate buffer (pH 6.0) for 10 min. Endogenous peroxidase activity was blocked with 0.3% hydrogen peroxide for 30 min. Sections were treated with rabbit VECTASTAIN Elite ABC horseradish peroxidase kit (Vector Laboratories, PK-6101) following the manufacturers protocols with all incubations were carried out at room temperature. Sections were developed in DAB (3,3′-diaminobenzidine) for 5 min, counterstained in haematoxylin, dehydrated and mounted in DPX. Staining was imaged on a Leica DM6 B using LASx. Primary antibodies: Cyclin T1 (AbCam, ab238940; 1:500), Myc (Y69; Abcam, ab32072, used at 1:1000) and Cleaved Caspase-3 (Asp175) (Cell Signalling technology, 9661; 1:400).

### Statistical analysis

Statistical analyses were performed using GraphPad Prism v9.0d (GraphPad Software, Inc., San Diego, CA, USA) as indicated with P ≤ 0.05 considered to be statistically significant.

### Data Availability

All sequencing datasets generated and used in this study, have been deposited in European Nucleotide Archive (www.ebi.ac.uk/ena) under accession code PRJEB52314. Proteomics data are available via ProteomeXchange with identifier PXD043114.

## Materials and Correspondence

Further information and requests for resources and reagents should be directed to, and will be fulfilled, by Dr Catherine Wilson.

## Author Contributions

CW, JH, GQ, AB conceptualised the study. AB, GQ, CB, EL, HR, CA, AS, VR, CW and MB performed the experimental work. KJ, QW, KH provided the modRNA AB performed the computation analysis assisted by JS. KS, BT and AS performed the live animal imaging. TK and AV assisted with animal work. KJ co-supervised AB. EP co-supervised GQ. SS co-funded and co-supervised AS. CW, JH, CA, AB, EL, GQ, CB, HR wrote or edited the manuscript. JH and CW obtained funding for the research.

## Acknowledgements

The authors thank the support staff in the Cambridge University Biomedical Services at the Anne McLaren Building and Cancer Research UK (CRUK) Cambridge Institute for 10x library prep and sequencing, and the QIMR Berghofer Medical Research Institute Proteomics facility for LC-MS acquisition. Some images were created with BioRender.com.

## Sources of funding

This work was supported by funding from the Wellcome Trust Institutional Strategic Support Fund (RG89529 & RG8930 to CHW), Academy of Medical Sciences Springboard award (G112756 to CHW), Rosetrees Trust Innovation and Translation Award (G106895 to CHW) and BHF project grant (G114642 to CHW). AB was funded by an AZ studentship (G108846 to CHW). EL was funded by a Cambridge BHF Centre of Research Excellence Studentship (RG96157). JH research is supported by grants and fellowships from the National Health and Medical Research Council of Australia and a Snow Medical Fellowship. SS was supported by a BHF Senior Fellowship (FS/18/46/33663).

## Declaration of Competing Interests

KJ, QW and KH are employees of AstraZeneca. MB and CHW are named inventors on a patent (PCT/GB2020/050350) relating to data in this manuscript.

