## Supplemental data for "A transient modified mRNA encoding Myc and Cyclin T1 induces cardiac regeneration and improves cardiac function after myocardial injury"

**Supplemental Figures 1 to 7**

**Supplemental Figure 1**

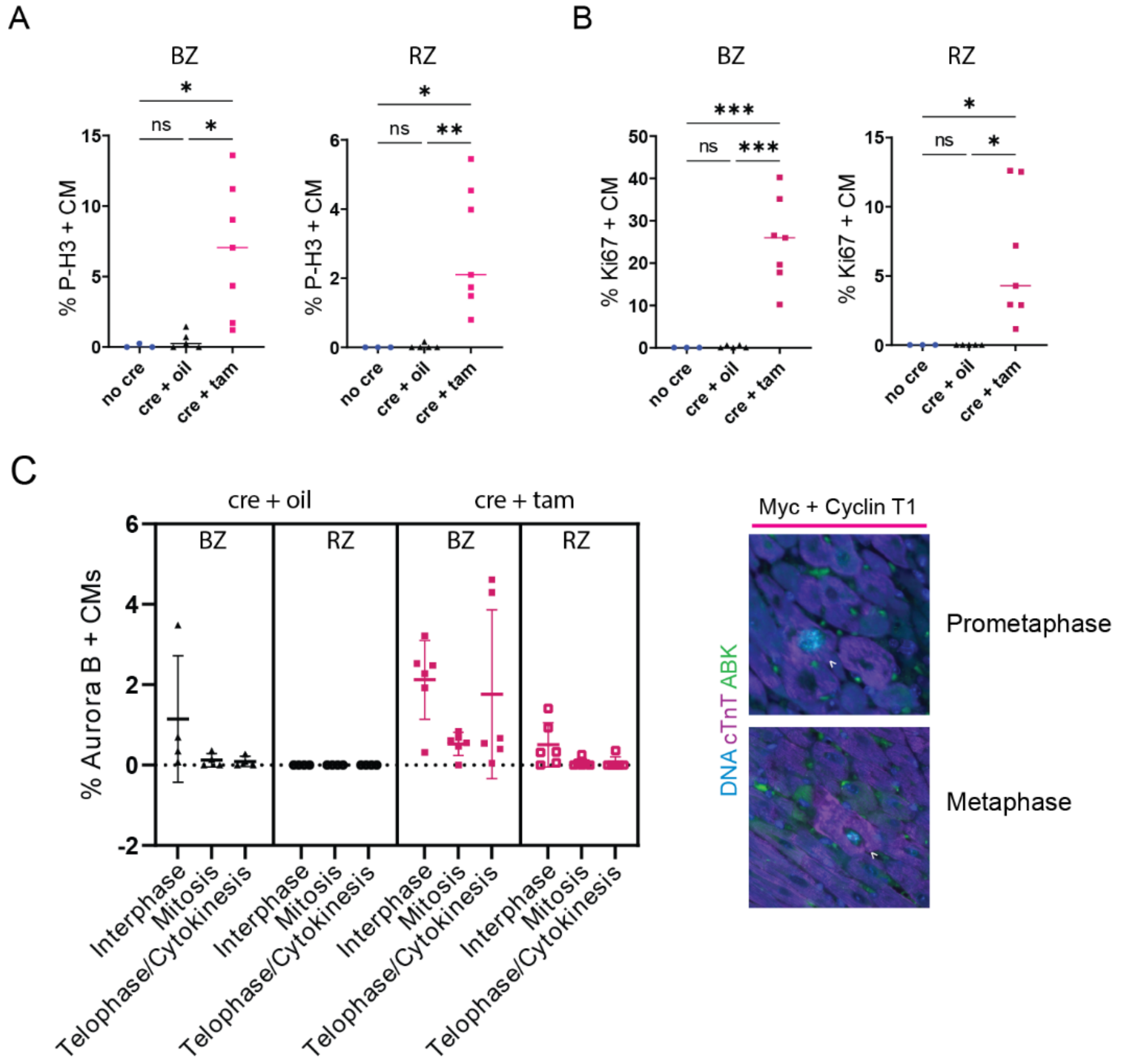


(A) Quantification of Immunofluorescence staining of P-H3 in the border or remote zone of MycER$^{T2}$ and Cyclin T1 (cre + tam, n = 7) and control (either oil treated (cre + oil, n = 5) or cre negative (no cre, n = 3)) hearts 48 hours following activation of MycER$^{T2}$ by tamoxifen treatment, as show in Figure 1A. One way-ANOVA with Tukey's multiple comparisons test; BZ, cre + tam to no cre p =

0.0313 (*), cre + tam to cre + oil p = 0.0176 (*). RZ, cre + tam to no cre p = 0.0154 (*), cre + tam to cre + oil p = 0.0058 (**). Mean shown.

(B) Quantification of Immunofluorescence staining of Ki67 in the border or remote zone of $MycER^{T2}$ and Cyclin T1 (cre + tam, n = 7) and control (either oil treated (cre + oil, n = 5) or cre negative (no cre, n = 3)) hearts 48 hours following activation of $MycER^{T2}$ by tamoxifen treatment, as show in Figure 1A. One way-ANOVA with Tukey's multiple comparisons test; BZ, cre + tam to no cre p = 0.0009 (***), cre + tam to cre + oil p = 0.0007 (***). RZ, cre + tam to no cre p = 0.0462 (*), cre + tam to cre + oil p = 0.0195 (*). Mean and SD shown.

(C) Immunofluorescence staining and breakdown of cell cycle stage of Aurora B Kinase (green), cardiac troponin (purple) and DNA (blue) in the border and remote zone of $MycER^{T2}$ and Cyclin T1 (cre + tam, n = 6) and control (either oil treated (cre + oil, n = 4)) hearts 48 hours following activation of $MycER^{T2}$ by tamoxifen treatment. Examples of prometaphase and metaphase ABK positive cardiomyocytes shown (white arrows).

#### Supplemental Figure 2

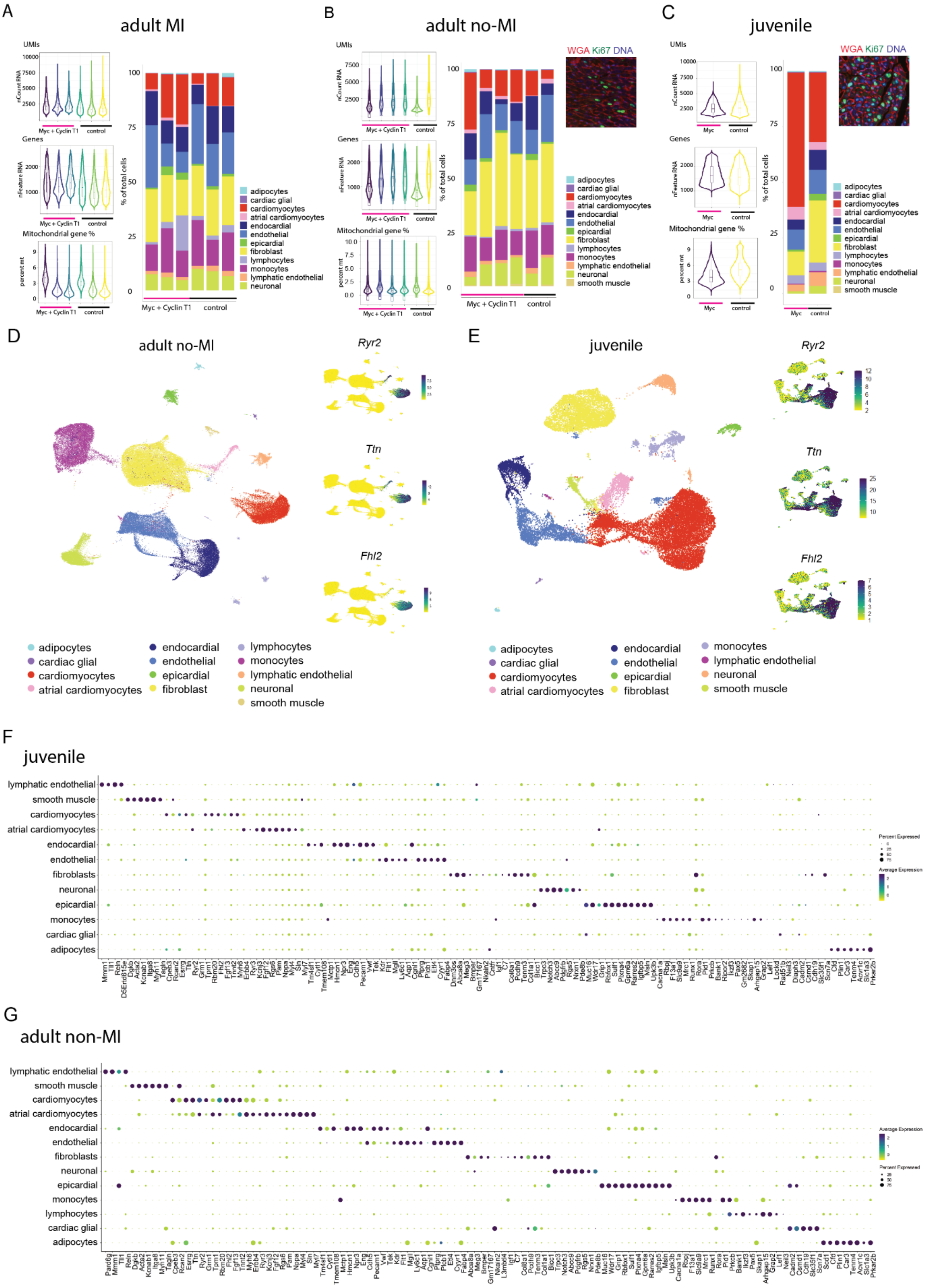

(A) Violin plots (left) displaying number of unique molecular identifiers (UMIs), Genes and mitochondrial gene % per nucleus for biological replicate samples in adult MI datasets. Bar plots (right) illustrating cell type abundance as % of total nuclei per sample.

(B) Violin plots and boxplots (left) displaying number of UMIs, Genes and mitochondrial gene % per nucleus for biological replicate samples in adult no-MI datasets. Bar plots (right) illustrating cell type abundance as % of total nuclei per sample. Representative immunofluorescence image of Ki67 (green) WGA (red) and DNA (blue) of adult no-MI MycER$^{T2}$ and Cyclin T1 expressing heart 48 hours following activation of MycER$^{T2}$ by tamoxifen treatment.

(C) Violin plot and boxplots (left) displaying number of UMIs, Genes and mitochondrial gene % per nucleus for biological replicate samples in juvenile datasets. Bar plots (right) illustrating cell type abundance as % of total nuclei per sample. For each nuclear sample, 3 hearts collected from mice of the same genotype were pooled together. Representative immunofluorescence image of Ki67 (green) WGA (red) and DNA (blue) of juvenile MycER$^{T2}$ expressing heart 48 hours following activation of MycER$^{T2}$ by tamoxifen treatment.

(D) UMAP plot (left) illustrating cell type clusters in adult no-MI dataset. UMAP feature plots (right) for cardiomyocyte specific markers *Ryr2*, *Ttn*, *Fhl2*.

(E) UMAP plot (left) illustrating cell type clusters in juvenile dataset. UMAP feature plots (right) for cardiomyocyte specific markers *Ryr2*, *Ttn*, *Fhl2*.

(F) Dot plot showing expression of cell type canonical markers per cell type cluster in adult no-MI dataset.

(G) Dot plot showing expression of cell type canonical markers per cell type cluster in juvenile dataset.

### Supplemental Figure 3

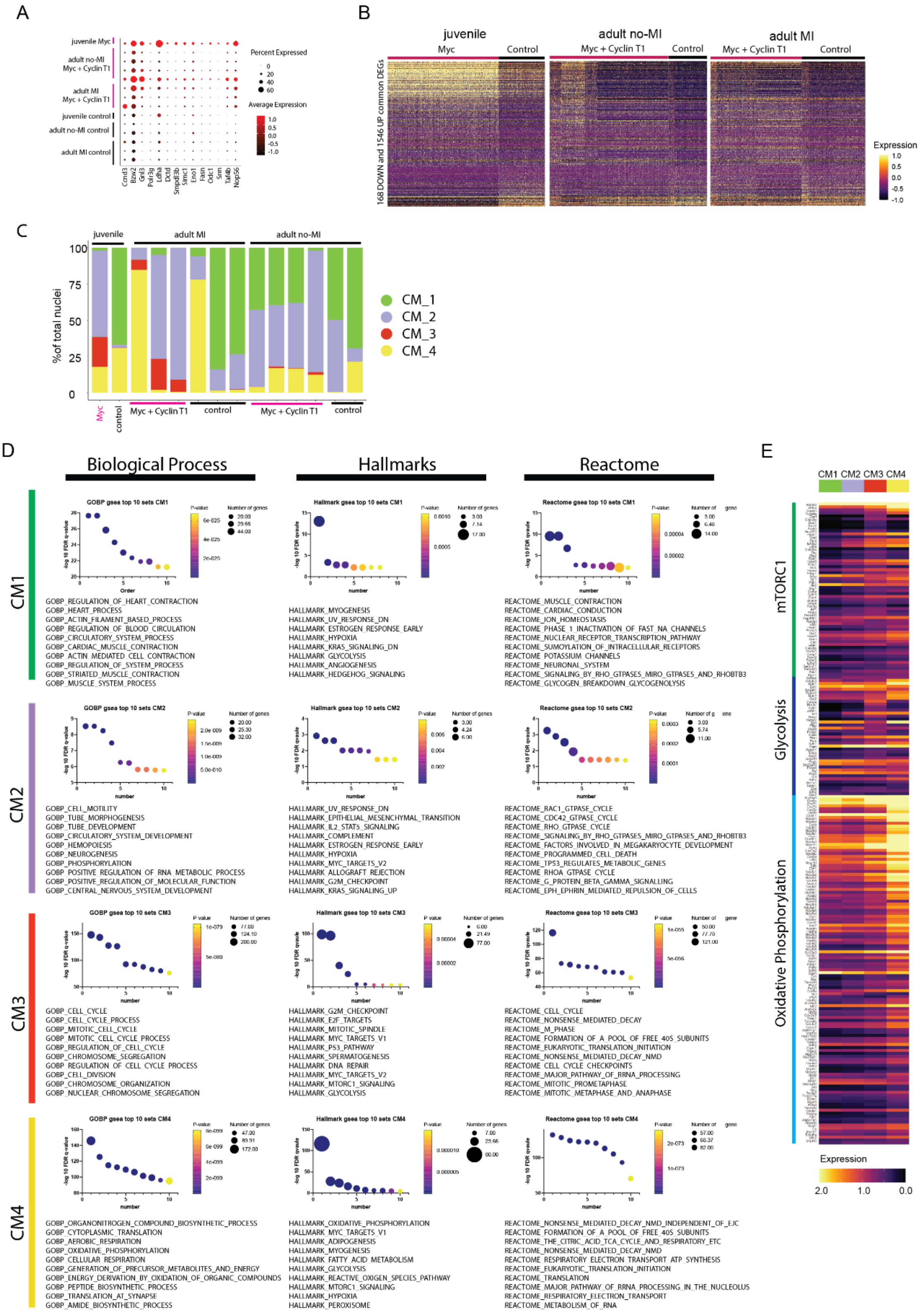

(A) Dot plot of common Myc targets grouped by biological sample and arranged by genotype.

(B) Heatmap for common upregulated and downregulated differentially expressed genes for each dataset grouped by genotype.

(C) Percent total nuclei for cardiomyocyte subcluster CM1, CM2, CM3, CM4 per biological replicate sample in adult no-MI, juvenile and adult MI datasets.

(D) Gene set enrichment analysis for cardiomyocyte subcluster CM1, CM2, CM3, CM4 markers (Wilcox test, $p_{adj} < 0.01$, $log2FC > 0.25$) for Gene Ontology biological process, Reactome and hallmark gene sets.

(E) Heatmap for differentially expressed genes for each gene set grouped by subcluster CM1, CM2, CM3, CM4 and for mTORC1 signalling, glycolysis and oxidative phosphorylation.

#### Supplemental Figure 4

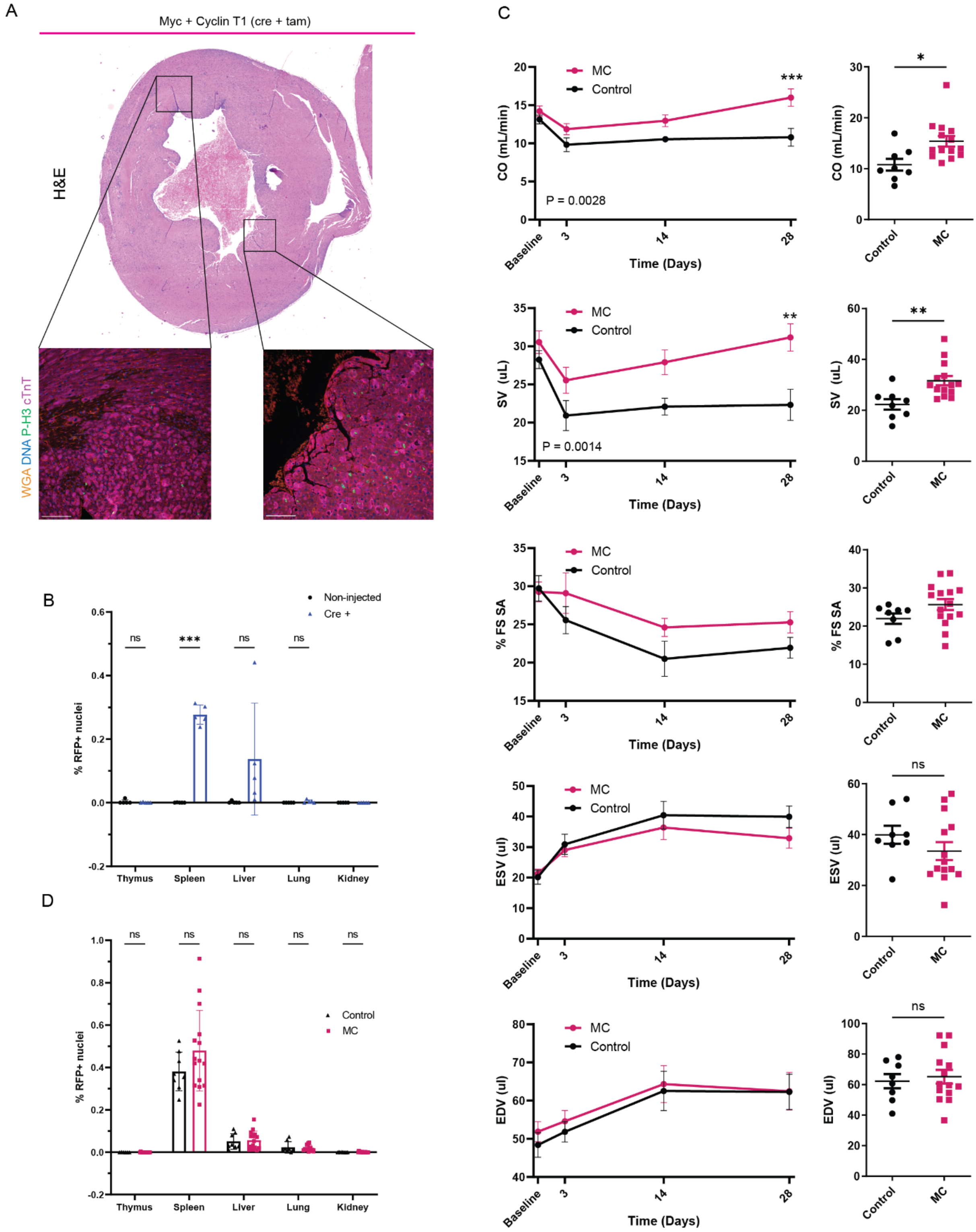


(A) H&E staining and immunofluorescence staining of P-H3 (green), cardiac troponin T (pink), WGA (orange) and DNA (Blue) in regions of MycER$^{T2}$ and

Cyclin T1 hearts following activation of $MycER^{T2}$ by tamoxifen treatment and at the time of culling when moribund (4 days post MI).

(B) Quantification of immunofluorescent staining for TdTomato positive nuclei, 5 days following intramyocardial injection with 100 µg Cre modRNA (n = 5) compared to untreated (n = 5) during surgery, immediately post-MI in thymus, spleen, liver, lung and kidney. Two way-ANOVA with Šídák's multiple comparisons non-injected vs Cre injected in the spleen p = 0.0002 (***). Mean and SD shown.

(C) Mean value and SEM over time (left) and 28 day comparison (right) of cardiac output (CO), stroke volume (SV), percent fractional shortening (% FS, short-axis), end systolic volume (ESV) and end diastolic volume (EDV) of MC (n = 12) and control mice (n = 9) over 28 days post MI. Mixed-effects model effect for CO vs treatment, p =0.0028; Šídák's multiple comparisons test p = 0.0005 at 28 days. Mixed-effects model effect for SV vs treatment, p = 0.0014; Šídák's multiple comparisons test p = 0.0029 at 28 days. Unpaired T-test at 28 days MC vs control mice, CO, p = 0.0110; SV, p = 0.0042.

(D) Quantification of immunofluorescent staining for TdTomato positive nuclei in $MycER^{T2}$ and Cyclin T1 (n = 14, MC) expressing mice and controls (n = 8), 28 days following intramyocardial injection with 100 µg Cre modRNA during surgery, immediately post-MI in thymus, spleen, liver, lung and kidney. Two-way ANOVA, ns = not significant. in 8 week old mice. Mean and SD shown.

#### Supplemental Figure 5

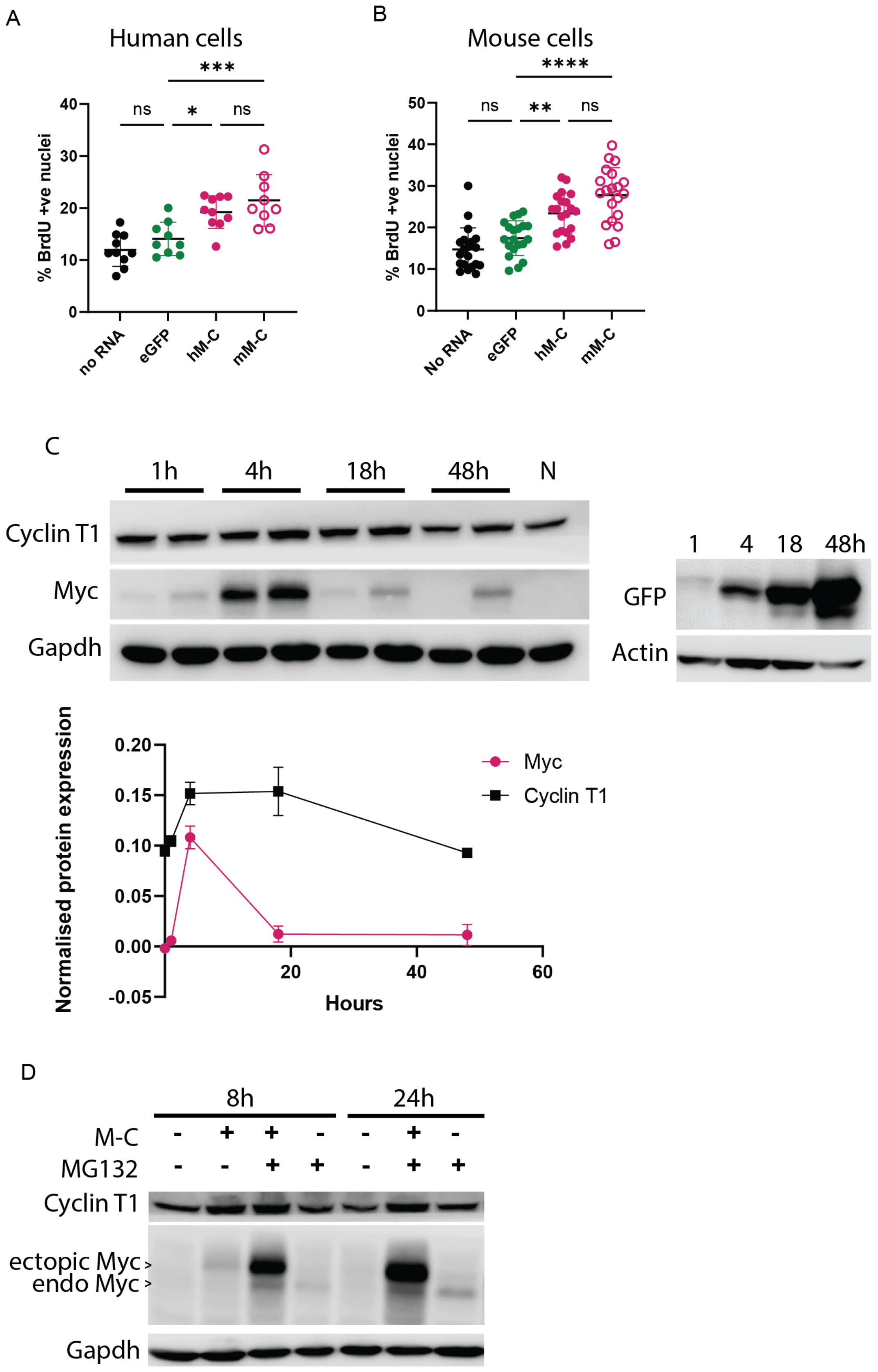

(A) Quantification of percent BrdU positive nuclei in lactate selected embryonically derived cardiomyocytes following treatment with eGFP, human (hM-C) or mouse (mM-C) Myc-Cyclin T1 modRNA. One-way ANOVA with Tukey's multiple comparisons test, eGFP vs hM-C, $p = 0.0218$; eGFP vs mM-C, $p = 0.0008$. Mean and SE shown.

(B) Quantification of percent BrdU positive nuclei in mouse embryo fibroblasts following treatment with eGFP, human (hM-C) or mouse (mM-C) Myc-Cyclin T1 modRNA. One-way ANOVA with Tukey's multiple comparisons test, eGFP vs hM-C, $p = 0.0054$; eGFP vs mM-C, $p < 0.0001$.

(C) Immunoblot (top) and quantification (bottom) of GFP, Myc and Cyclin T1 protein levels in lactate selected embryonically derived cardiomyocytes following mM-C treatment at 1-, 4-, 18- and 48-hours post treatment. Mean and SEM shown.

(D) Immunoblot of Myc and Cyclin T1 protein levels in lactate selected embryonically derived cardiomyocytes following mM-C treatment together with protease inhibitor MG132 at 8- and 24-hours post treatment.

#### Supplemental Figure 6

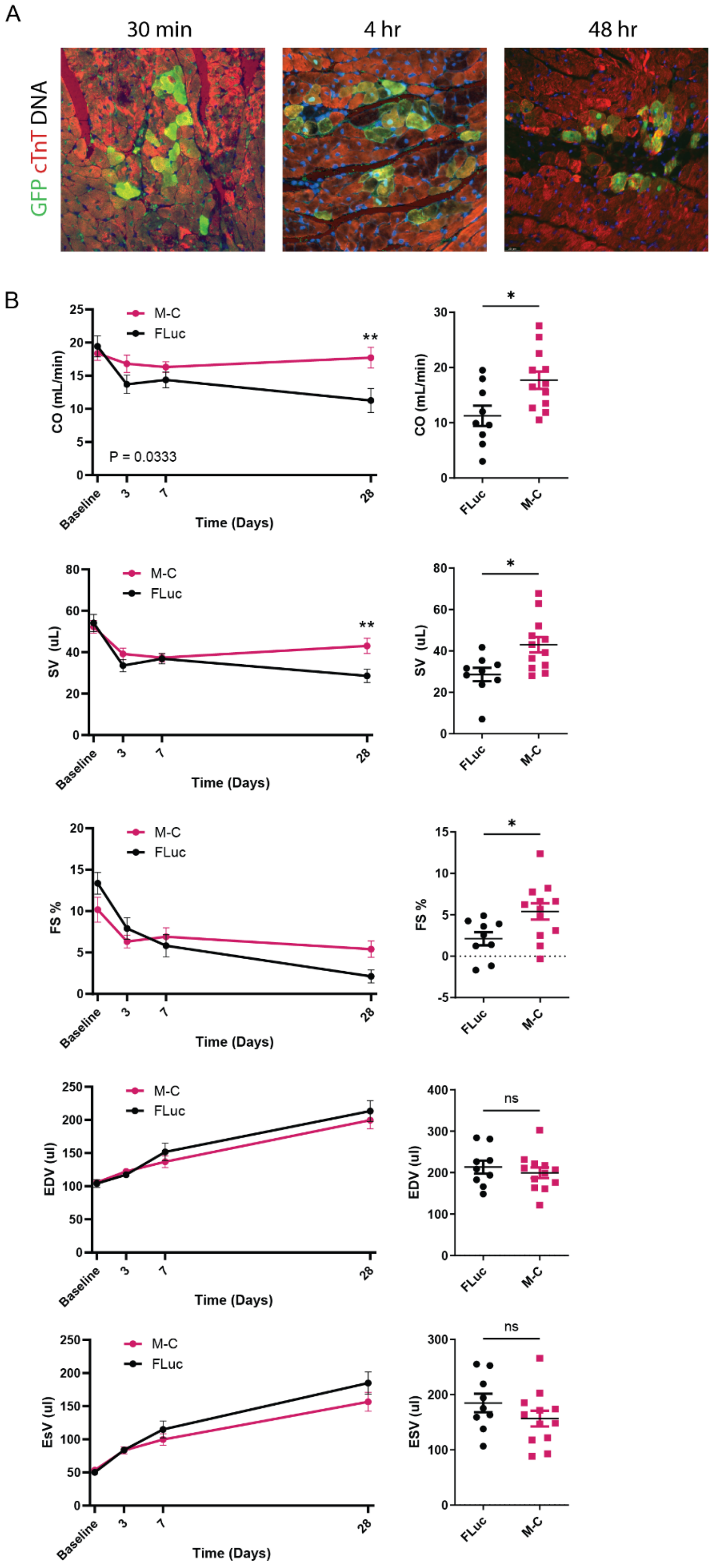

(A) Immunofluorescence staining of GFP (green), cardiac troponin (cTnT, red) and DNA (blue) indicating cardiac expression of GFP at 30 minutes, 4 hours, and 48 hours post GFP modRNA injection.

(B) Mean value and SEM over time (left) and 28 day comparison (right) of cardiac output (CO), stroke volume (SV), percent fractional shortening (% FS), end systolic volume (ESV) and end diastolic volume (EDV) of M-C and FLuc treated mice over 28 days post MI. Mixed-effects model effect for CO vs treatment, $p = 0.0333$; Šídák's multiple comparisons test $p = 0.0051$ at 28 days. Mixed-effects model effect for SV vs treatment, Šídák's multiple comparisons test $p = 0.0072$ at 28 days. Unpaired T-test at 28 days MC vs control mice, CO, $p = 0.0148$; SV, $p = 0.0110$; %FS, $p = 0.0233$.
